# University-wide chronotyping shows late-type students have lower grades, shorter sleep, poorer well-being, lower self-regulation, and more absenteeism

**DOI:** 10.1101/2021.08.04.455177

**Authors:** Sing Chen Yeo, Jacinda Tan, Clin K.Y. Lai, Samantha Lim, Yuvan Chandramoghan, Fun Man Fung, Patricia Chen, Timothy J. Strauman, Joshua J. Gooley

## Abstract

A person’s preferred timing of nocturnal sleep (chronotype) has important implications for cognitive performance. Students who prefer to sleep late may have a selective learning disadvantage for morning classes due to inadequate sleep and circadian desynchrony. Here, (1) we tested whether late-type students perform worse only for morning classes, and (2) we investigated factors that may contribute to their poorer academic achievement. Chronotype was determined objectively in 33,645 university students (early, *n*=3,965; intermediate, *n*=23,787; late, *n*=5,893) by analyzing the diurnal distribution of their logins on the university’s Learning Management System (LMS). Late-type students had lower grades than their peers for courses held at all different times of day, and during semesters when they had no morning classes. Actigraphy studies (*n*=261) confirmed LMS-derived chronotype was associated with students’ sleep patterns. Nocturnal sleep on school days was shortest in late-type students because they went to bed the latest and woke up early compared with non-school days. Surveys showed that late-type students had lower self-rated health and mood (*n*=357), and lower metacognitive self-regulation (*n*=752). Wi-Fi connection data for classrooms (*n*=17,356) revealed that late-type students had lower lecture attendance than their peers for classes held in both the morning and the afternoon. Our findings suggest that multiple factors converge to impair learning in late-type students. Shifting classes later can improve sleep and circadian synchrony in late-type students but is unlikely to eliminate the performance gap. Interventions that focus on improving students’ well-being and learning strategies may be important for addressing the late-type academic disadvantage.

## Introduction

The timing of nocturnal sleep has important implications for students’ academic achievement. There are individual differences in diurnal preferences for waking activities and sleep (“chronotype”) (1), whereby some students have an earlier or later sleep-wake pattern than others. These differences are thought to be driven by biological and environmental factors that converge to influence students’ sleep behavior (2). Studies in children, adolescents, and undergraduates have shown that late-type students (“evening types” or “owls”) have lower grades compared with early-type students (“morning types” or “larks”) (1, 3). Meta-analyses indicate that later chronotype (i.e., greater eveningness or lower morningness) is associated with a small negative effect on academic achievement in students attending high schools and universities (4). Given the influence of grades in determining post-graduate career opportunities (5), it is important to characterize factors that contribute to poorer academic achievement in late-type students. Doing so will pave the way for interventions to minimize the academic disadvantage associated with later chronotype.

Previous studies have identified several mechanisms that could explain the association between chronotype and academic performance. Late-type students with morning classes have shorter nocturnal sleep than their peers because they have later bedtimes but wake up at about the same time (6–9). Late-type students may also have higher circadian drive for sleep during morning classes due to a later phase of circadian entrainment (10, 11). The combined influence of insufficient sleep and circadian desynchrony can give rise to daytime sleepiness and impaired cognitive performance (12). Later chronotype is also associated with poorer self-rated health (13, 14), sleep problems (6, 15), mood disturbances and psychological disorders (16, 17), and depression symptoms (18), all of which could impact students’ performance at school. Psychological correlates of learning are also lower in students with later chronotype, including self-regulation (19, 20), self-control (21, 22), intrinsic motivation (23, 24), learning goals (25, 26), conscientiousness (24, 27, 28), and grit (29). Lastly, late-type students may have lower class attendance rates, e.g. due to oversleeping or feeling tired and unmotivated (6). It is likely that multiple factors combine to impair performance in late-type students. However, the relative importance of these factors is difficult to interpret due to differences across studies in the research design (e.g., the way that chronotype was defined) and outcome measures that were evaluated (e.g., few studies examined more than one outcome).

It is widely assumed that late-type students have an academic disadvantage for classes and examinations held in the morning, but not later in the day (1). While some studies support this assumption (9, 30–33), others found that later chronotype was associated with poorer performance in both the morning and the afternoon (34–37). However, there are several methodological issues that make it difficult to interpret previous findings. Some studies did not test for an interaction between chronotype and time of day in their statistical models (30, 32, 34, 35). Other studies excluded intermediate-type students from the analysis (34, 35), or collapsed performance data across multiple class start times in ways that were either not necessary or not clearly justified (31, 33–35). It is also uncertain if results from high schools where students attended classes only in the morning or only in the afternoon/evening (9) are generalizable to universities where students’ timetables differ across the week. All studies included convenience samples with chronotype determined by self-report, which may have resulted in response-bias, self-selection bias, and self-reporting bias. Moreover, it is unclear whether previous findings that reached statistical significance were meaningful in terms of effect size. Given the limitations associated with prior work, additional studies are needed to clarify the relationship between chronotype and time of day on academic achievement in university students.

Recent work suggests that chronotype can be determined objectively by analyzing the temporal distribution of students’ logins on the university’s Learning Management System (LMS) (37). The key advantage of using LMS logins for chronotyping is that data are collected passively at large scale, making it possible to measure chronotype in large samples of students while minimizing sources of bias. This method was used at a university in the American Midwest to show that late-type students (“owls”) had significantly lower academic performance than early-type students (“larks”) and intermediate-type students (“finches”) for courses held in the morning, afternoon, and evening (37). These findings challenged the view that shifting classes and exams to the afternoon/evening can reduce or eliminate the performance gap in late-type students. Given the strong theoretical basis for late-type students having a greater academic disadvantage for morning classes, it is important to test whether these results can be reproduced and extended in other universities and sociocultural contexts. Importantly, the LMS chronotyping method has not been validated by measuring students’ sleep patterns. While students must be awake to log in to the LMS, their diurnal pattern of LMS usage may be influenced by environmental/social factors unrelated to sleep. Additionally, it has not been tested whether LMS-derived chronotype associates with factors that are important for students’ learning including sleep health, well-being, psychological characteristics, and class attendance. Addressing these knowledge gaps will enable universities to develop targeted solutions to improve learning in late-type students.

The objective of our study was to use the LMS chronotyping method to test whether late-type students have an academic disadvantage. University-wide chronotyping was performed to test associations of chronotype with academic performance and learning-related behaviors. First, we tested the hypothesis that late-type students have a lower grade point average than their peers for courses held in the morning but not in the afternoon/evening. Second, we tested the hypothesis that later chronotype is associated with a later nocturnal sleep pattern and shorter sleep duration on days with morning classes. Third, we tested the hypothesis that late-type students fare more poorly on indices of well-being, including sleep quality, daytime sleepiness, self-rated health, and mood. Fourth, we tested the hypothesis that late-type students have lower scores for psychological correlates of learning, including self-regulation, self-control, intrinsic motivation, learning goals, conscientiousness, and grit. Fifth, we tested the hypothesis that late-type students have lower attendance rates for lecture courses.

## Results

### Categorization of students into chronotype groups

The diurnal time courses of LMS logins were examined separately on non-school days and school days in 33,645 students (83% of all students enrolled in at least one course) whose data were compiled over 5 semesters (Fig. 1a). In each student, the temporal distribution of LMS logins was used to derive his/her median login phase of activity (Fig. 1b). The population distribution of median login phase on non-school days was then used to assign students to chronotype categories (Fig. 1c). LMS-derived chronotype was determined using data for non-school days because students are more likely to follow their preferred sleep-wake pattern when they do not have classes. Individuals whose login phase was more than one standard deviation earlier or later than the population median were defined as early-type students (11.8%, *n* = 3,965) and late-type students (17.5%, *n* = 5,893), respectively (37). The remaining individuals were defined as intermediate-type students (70.7%, *n* = 23,787). Chronotype groups did not differ by sex and there were minor differences for ethnicity and country of citizenship/residency (Supplementary Table 1). However, early-type students were about a year older than their peers with an earlier date of matriculation (i.e., they were closer to graduating), and they were more likely to live off campus. Late-type students matriculated more recently and were more likely to live in residence halls or colleges on campus. There was some variation in chronotype across different schools within the university, in which the School of Continuing and Lifelong Education (a school which caters to working adults) had a greater proportion of early-type students, and the Yong Siew Toh Conservatory of Music had a greater proportion of late-type students (Supplementary Table 1).

**Fig. 1.**
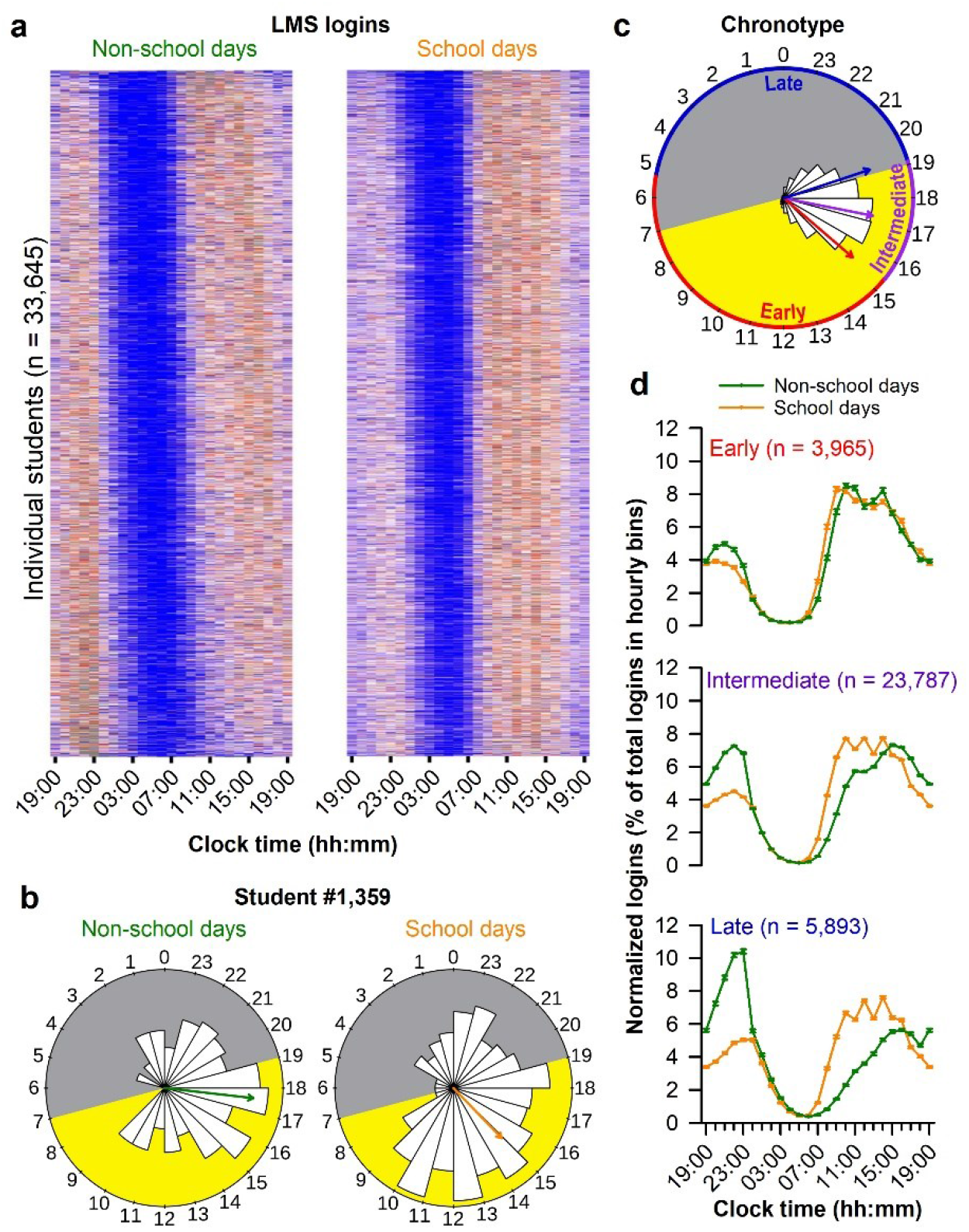
Learning Management System (LMS)-derived chronotype. (**a**) The heat maps show diurnal time courses of LMS logins in 33,645 students on non-school days and school days. High and low numbers of logins are indicated by red and blue, respectively. Each row corresponds to an individual student whose logins were binned hourly. Data were compiled over 5 semesters and normalized by expressing logins in each 1-h bin as a percentage of the total logins in each student. (**b**) The circular histograms show the number of LMS logins at different times of day in a representative student (Student #1,359). The yellow and grey shading indicates the approximate hours of daylight and darkness, respectively, where the data were collected (Singapore, 1.35°N of the equator). The arrows indicate the median login phase determined using circular statistics. (**c**) The circular histogram shows the population distribution of median login phase on non-school days, which was used to categorize students into chronotype groups. Students whose login phase was more than one standard deviation earlier (red arrow) or later (blue arrow) than the population median (purple arrow) were defined as early-type students and late-type students, respectively. (**d**) The diurnal rhythm (mean ± 95% CI) of LMS logins is shown for non-school days and school days in each chronotype group.

Next, we compared the diurnal time courses of login behavior between non-school days and school days for the different chronotype groups (Fig. 1d). In each student, we measured the difference in his/her median login phase on non-school days and school days to derive a relative measure of social jet lag (37). Early-type students exhibited a diurnal rhythm of LMS logins that was only slightly later for non-school days compared with school days (Social jet lag = 0.32 h, 95% CI = 0.26 to 0.37 h). In contrast, intermediate-type students exhibited a substantially later rhythm of LMS logins on non-school days relative to school days (Social jet lag = 2.28 h, 95% CI = 2.26 to 2.30 h). Late-type students showed the greatest temporal discrepancy in login behavior (Social jet lag = 4.69 h, 95% CI = 4.62 to 4.76 h), with a later and more gradual rise of LMS activity on non-school days relative to the other chronotype groups (Fig. 1d).

### Chronotype and grade point average

LMS-derived chronotype was significantly associated with cumulative grade point average (*F*_2,33610_ = 204.7, *P* < 0.001; Fig. 2a), adjusting for sex, age, ethnicity, country of citizenship/residency, year of matriculation, school of enrollment, and type of residence. Early-type students performed marginally better than intermediate-type students (Tukey test, *P* < 0.001; Cohen’s d = 0.05, 95% CI: 0.02 to 0.09). By comparison, grade point average was substantially lower in late-type students compared with intermediate-type students (Tukey test, *P* < 0.001; Cohen’s d = −0.27, 95% CI: −0.30 to −0.24).

**Fig. 2.**
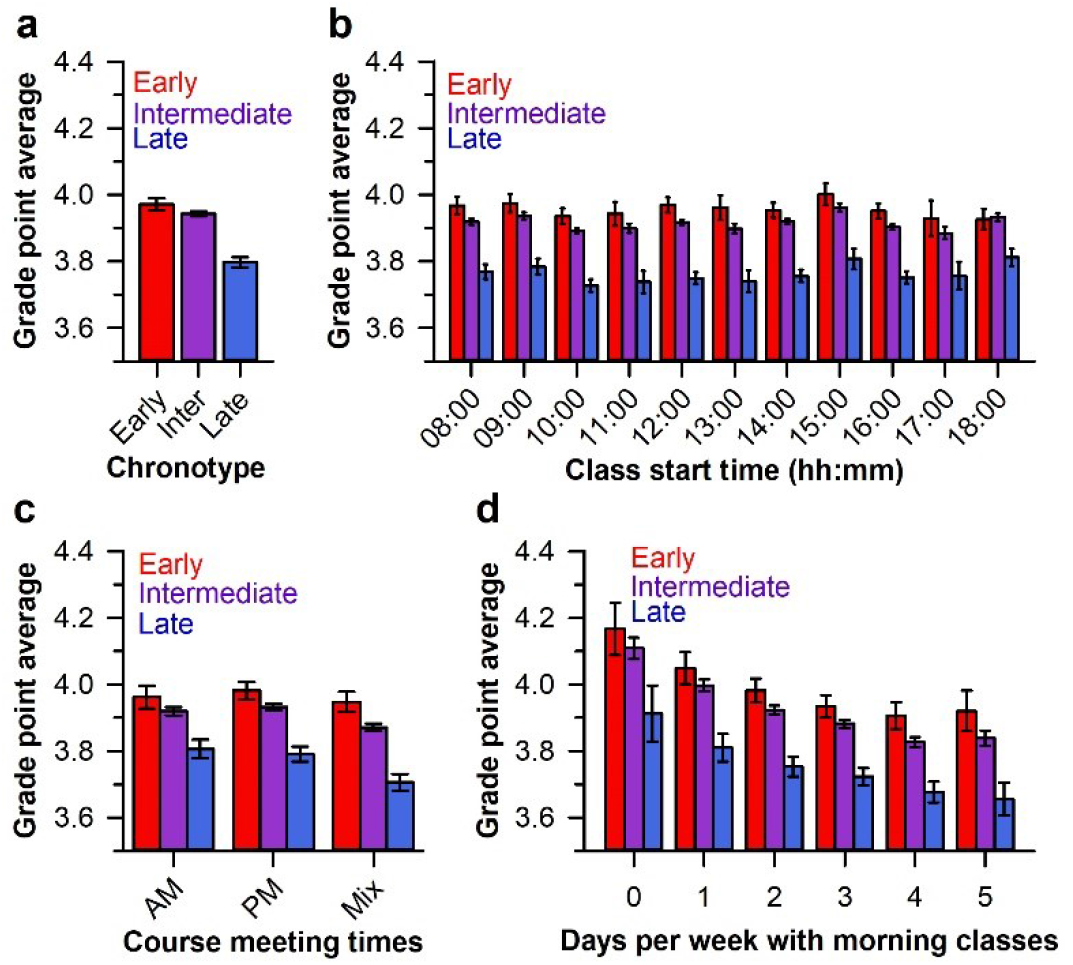
Late-type students had lower academic performance. Grade point average (GPA) was compared between students who were categorized into chronotype groups (early, *n* = 3,965; intermediate, *n* = 23,787; late, *n* = 5,893) using their Learning Management System login data. Results are shown for (**a**) cumulative GPA, (**b**) GPA sorted by the primary meeting time of a given course (e.g., lecture or seminar), (**c**) GPA sorted by the timing of all class components within a course (AM, all classes started before 12:00; PM, all classes started at 12:00 or later; Mix, classes took place in both the morning and the afternoon/evening), and (**d**) GPA sorted by the number of days of the week that students had morning classes. In all analyses, late-type students had a lower GPA compared with the other chronotype groups. The mean ± 95% CI is shown in each bar graph.

Next, we conducted a set of 3 analyses to test whether the association of chronotype with academic performance was modulated by class start time (Fig. 2b-d; Supplementary Fig. 1), adjusting for covariates. In our first analysis of time-of-day effects, class start time was defined as the primary meeting time of a given course (e.g., lecture or seminar). There was a significant interaction between chronotype and class start time (*F*_20, 213350_ = 2.0, *P* = 0.004), but the difference in grades between chronotype groups did not vary systematically by time of day (Fig. 2b; Supplementary Fig. 1). Multiple comparison tests for all class start times ranging from 08:00 to 18:00 showed that early-type students performed marginally better than intermediate-type students for classes at 12:00 (Tukey test, *P* = 0.02), but grade point average did not differ between these groups for any other class start time (Tukey test, *P* > 0.06 for all pairwise comparisons; Cohen’s d range = −0.01 to 0.08). In contrast, grade point average was lower in late-type students for all class start times relative to intermediate-type students (Tukey test, *P* < 0.001 for all pairwise comparisons; Cohen’s d range: −0.27 to −0.15) (Fig. 2b; Supplementary Fig. 1).

In our dataset, it was common for individual courses to have classes (e.g., lectures, tutorials, and laboratories) that met at different times of the day. In our second analysis of time-of-day effects, we therefore categorized courses as occurring only in the morning (all classes starting before 12:00), only in the afternoon/evening (all classes starting at 12:00 or later), or in both the morning and the afternoon/evening (Fig. 2c). There was a significant interaction between chronotype and course time on grades (*F*_4,36199_ = 3.9, *P* = 0.004). Multiple comparison tests showed that early-type students performed slightly better than intermediate-type students for courses with mixed morning/afternoon/evening start times (Tukey test, *P* < 0.001; Cohen’s d = 0.11, 95% CI: 0.06 to 0.16). For all course times, late-type students had a lower grade point average compared with intermediate-type students (Tukey test, *P* < 0.001 for all pairwise comparisons; Cohen’s d range: −0.23 to −0.16) (Fig. 2c; Supplementary Fig. 1).

In our third analysis of time-of-day effects, we tested whether the association of chronotype with grade point average varied by the number of days of the week that students had morning classes in each semester (Fig. 2d). There was no interaction of chronotype and the number of days with morning classes on grade point average (*F*_10, 41866_ = 1.0, *P* = 0.44). Main effects showed that later chronotype (*F*_2, 33544_ = 144.1, *P* < 0.001) and more days per week with morning classes (*F*_5, 41496_ = 46.5, *P* < 0.001) were associated with poorer academic performance. The effect size of being a late-type student on grade point average (Cohen’s d range: −0.35 to - 0.24) was greater than the effect size of being an early-type student (Cohen’s d range: 0.06 to 0.13) (Supplementary Fig. 1). Notably, late-type students had lower academic performance than intermediate-type students even when they never had morning classes (Cohen’s d = −0.35, 95% CI: −0.51 to −0.20) (Fig. 2d; Supplementary Fig. 1).

### Chronotype and sleep behavior

Self-reported sleep behavior was investigated in 357 students who completed surveys and were categorized by their LMS-derived chronotype (early, *n* = 24; intermediate, *n* = 252; late, *n* = 81) (Fig. 3a; Supplementary Table 2). There was a significant interaction between chronotype and type of day (non-school day, school day) on self-reported wake-up time, midpoint of time in bed (TIB), and nocturnal sleep duration, but not bedtime (Supplementary Table 3), adjusting for sex, age, ethnicity, country of citizenship/residency, class year, school of enrollment, and type of residence. Main effects indicated that later chronotype was associated with later and shorter nocturnal sleep, and non-school days were associated with later and longer nocturnal sleep (Fig. 3a; Supplementary Tables 2 and 3). On non-school days, bedtime and wake-up time were almost an hour earlier in early-type students compared with intermediate-type students. Conversely, bedtime and wake-up time were nearly an hour later in late-type students (Fig. 3a; Supplementary Table 4). Multiple comparison tests showed that wake-up time and midpoint of TIB differed between all chronotype groups on non-school days (Tukey’s test, *P* < 0.025 for all pairwise comparisons), but there were no differences between chronotype groups for self-reported nocturnal sleep duration (Tukey’s test, *P* ≥ 0.99 for all pairwise comparisons) (Fig. 3a; Supplementary Table 4).

**Fig. 3.**
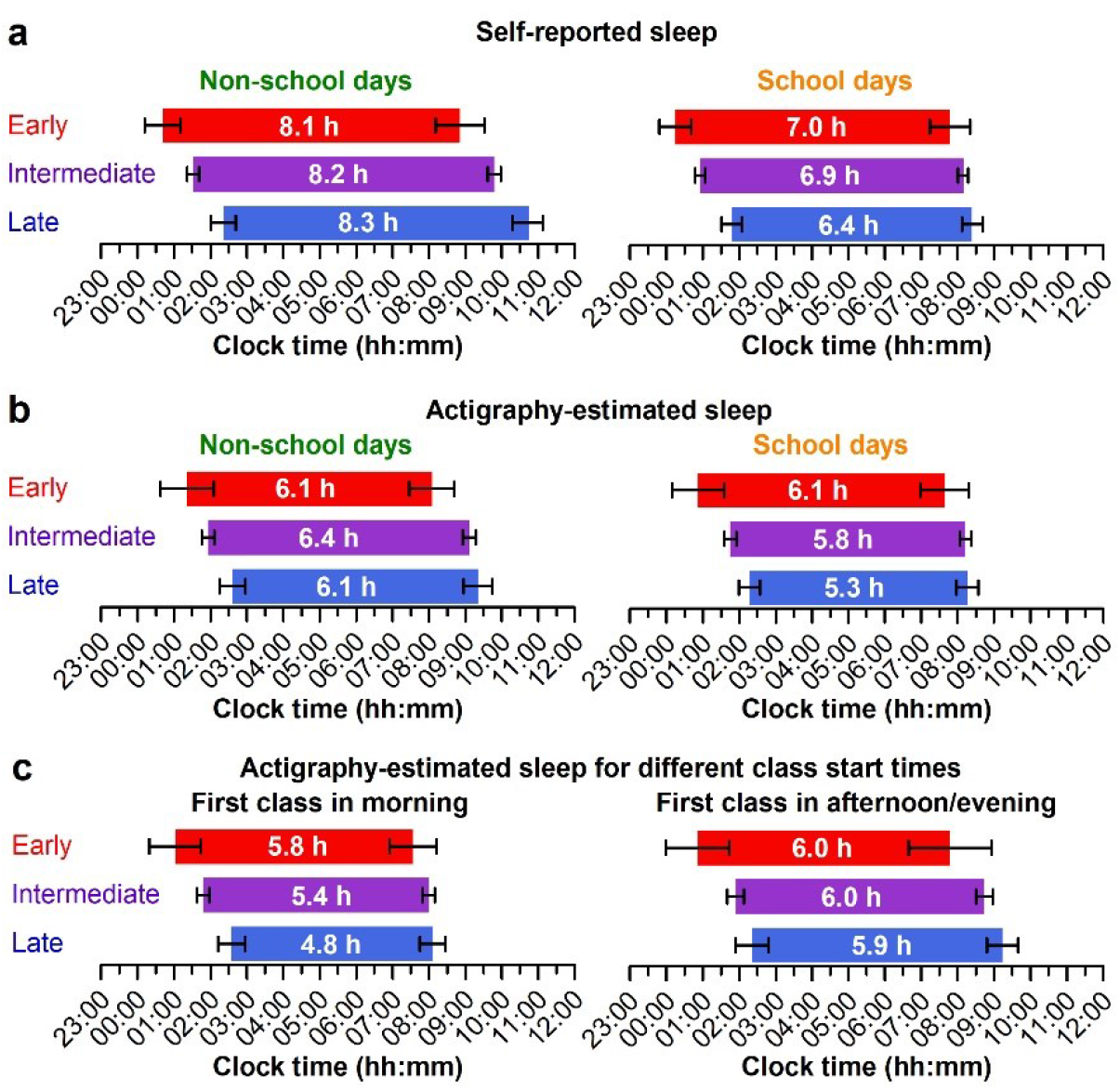
Late-type students had shorter sleep on school days. Nocturnal sleep behavior on non-school days and school days was compared between students who were categorized into chronotype groups using their Learning Management System login data. (**a**) Self-reported sleep was assessed using surveys (Chronotype: early, *n* = 24; intermediate, *n* = 252; late, *n* = 81) and (**b**) objective sleep behavior was estimated using actigraphy watches (Chronotype: early, *n* = 18; intermediate, *n* = 189; late, *n* = 54). (**c**) Actigraphy-estimated sleep periods on school days were sorted by days that students’ first class took place in the morning versus the afternoon/evening. In panel **a**, each horizontal bar indicates the time-in-bed interval for sleep. The mean ± 95% CI is shown for self-reported bedtime and wake-up time, and the mean sleep duration is shown in each bar. In panels **b** and **c**, each bar indicates the actigraphy-estimated sleep period. The mean ± 95% CI is shown for sleep onset and sleep offset, and the total sleep time is shown in each bar.

On school days, self-reported bedtime was about 40 minutes earlier in early-type students and about 50 minutes later in late-type students compared with intermediate-type students (Fig. 3a; Supplementary Table 4). However, wake-up times did not differ significantly between chronotype groups (Tukey’s test, *P* > 0.78 for all pairwise comparisons). In late-type students, the midpoint of TIB on school days was significantly later (Tukey’s test, *P* = 0.023; Cohen’s d = 0.56, 95% CI: 0.29 to −0.85) and nocturnal sleep duration was about 30 minutes shorter (Tukey’s test, *P* = 0.003; Cohen’s d = −0.44, 95% CI: −0.74 to −0.17) compared with intermediate-type students (Supplementary Table 4). Later chronotype was also associated with a greater difference in the midpoint of sleep between school days and non-school days (*F*_2, 341_ = 6.1, *P* = 0.002), in which social jet lag was 22 minutes greater in late-type students relative to intermediate-type students (Tukey’s test, *P* = 0.007; Cohen’s d = 0.35, 95% CI: 0.06 to 0.61).

Next, we compared objective sleep behavior in a subset of 261 students who wore an actigraphy watch for 10-42 days (Chronotype: early, *n* = 18; intermediate, *n* = 189; late, *n* = 54) (Fig. 3b; Supplementary Table 2). There was a significant interaction between chronotype and type of day on sleep offset and nocturnal total sleep time, but not sleep onset or midpoint of sleep (Supplementary Table 3), adjusting for covariates. Main effects showed that later chronotype was associated with later and shorter nocturnal sleep, and non-school days were associated with later and longer sleep (Fig. 3b; Supplementary Tables 2 and 3). On non-school days, early-type students fell asleep about 30 minutes earlier than intermediate-type students, and they woke up about an hour earlier (Tukey’s test, *P* = 0.004) (Fig. 3b; Supplementary Table 4). By comparison, late-type students fell asleep about 30 minutes later but woke up at about the same time as intermediate-type students. Consequently, the midpoint of sleep was about 45 minutes earlier in early-type students and nearly 30 minutes later in late-type students. However, nocturnal total sleep time on non-school days did not differ between chronotype groups (Tukey’s test, *P* > 0.45 for all pairwise comparisons) (Fig. 3b; Supplementary Table 4).

On school days, early-type students fell asleep about 50 minutes earlier and woke up about 35 minutes earlier than intermediate-type students. Late-type students fell asleep about 30 minutes later but woke up at about the same time as intermediate-type students. Multiple comparison tests did not detect a difference in sleep offset on school days between chronotype groups (Tukey’s test, *P* > 0.38 for all pairwise comparisons). The midpoint of sleep was about 45 minutes earlier in early-type students and about 20 minutes later in late-type students. Multiple comparison tests showed that nocturnal total sleep time in late-type students was about 30 minutes shorter on school days compared with intermediate-type students (Tukey’s test, *P* = 0.028) (Fig. 3b; Supplementary Table 4). Chronotype did not associate with the magnitude of social jet lag between school days and non-school days (*F*_2, 235_ = 2.0, *P* = 0.13).

Next, we investigated whether associations of chronotype with sleep behavior varied by class start time (Fig. 3c; Supplementary Table 5). We compared actigraphy-estimated sleep between chronotype groups when students’ first class of the day took place in the morning (starting before 12:00) versus in the afternoon/evening (starting at 12:00 or later). There was a significant interaction between chronotype and class start time for nocturnal total sleep time (*F*_2,235_ = 3.8, *P* = 0.024), but not for sleep onset (*F*_2,235_ = 1.3, *P* = 0.28), sleep offset (*F*_2,231_ = 2.3, *P* = 0.10) or midpoint of sleep (*F*_2,231_ = 0.62, *P* = 0.54). Main effects showed that later chronotype was associated with later sleep onset (*F*_2,245_ = 8.3, *P* < 0.001), later midpoint of sleep (*F*_2,243_ = 6.1, *P* = 0.003), and shorter nocturnal sleep (*F*_2,241_ = 4.8, *P* = 0.009), whereas sleep offset did not differ significantly between groups (*F*_2,240_ = 3.0, *P* = 0.051). Morning classes were associated with an earlier sleep offset (*F*_1,232_ = 30.1, *P* < 0.001), earlier midpoint of sleep (*F*_1,232_ = 8.2, *P* = 0.004), and shorter nocturnal sleep (*F*_1,237_ = 32.0, *P* < 0.001); however, there was no main effect of class start time on sleep onset (*F*_1,235_ = 0.53, *P* = 0.47). Late-type students and intermediate-type students obtained significantly less nocturnal sleep on days when their first class was in the morning rather than in the afternoon/evening (about 60 minutes and 40 minutes, respectively; Tukey’s test, *P* < 0.001 for both pairwise comparisons). In contrast, sleep duration in early-type students did not differ significantly for morning and afternoon/evening class start times (Tukey’s test, *P* = 0.92). Multiple comparison tests showed that late-type students had shorter nocturnal sleep than their peers when they had morning classes (Tukey’s test, *P* < 0.008 for both pairwise comparisons), whereas there were no differences in sleep duration between chronotype groups when students’ first class of the day took place in the afternoon/evening (Tukey’s test, *P* > 0.90 for all pairwise comparisons). On days with morning classes, late-type students had about 40 minutes less sleep than intermediate-type students (Cohen’s d = −0.67, 95% CI: −0.98 to −0.34) (Supplementary Table 5).

### Chronotype, well-being, and psychological correlates of learning

Survey data were used to test for differences in well-being between students who were categorized by their LMS-derived chronotype (early, *n* = 24; intermediate, *n* = 252; late, *n* = 81). Late-type students fared more poorly on well-being measures relative to the other chronotype groups (Fig. 4a; Supplementary Table 6). Kruskal-Wallis tests showed that chronotype was associated with self-rated health (*H* = 7.1, *P* = 0.029), and feelings of fatigue/ low motivation (*H* = 12.0, *P* = 0.003) over the past week. Multiple comparison tests showed that self-rated health and fatigue/ low motivation were significantly worse in late-type students compared with intermediate-type students (Dunn’s test, *P* < 0.05 for both comparisons), but did not differ between early-type students and intermediate-type students (Dunn’s test, *P* > 0.35 for both comparisons). Differences between chronotype groups for sleep quality, daytime sleepiness, sadness, lack of focus, and anxiety did not reach statistical significance (*H* ≤ 5.0, *P* > 0.08 for all measures). Effect sizes were small for late-type students for self-rated health (Cliff’s delta = 0.16, 95% CI: 0.03 to 0.29), fatigue/ low motivation (Cliff’s delta = 0.21, 95% CI: 0.07 to 0.34), and lack of focus (Cliff’s delta = 0.15, 95% CI: 0.01 to 0.28) relative to intermediate-type students, and the 95% CIs overlapped marginally with zero for the other well-being measures (Fig. 4a; Supplementary Table 7). Effect sizes were negligible for early-type students for all well-being measures (Cliff’s delta range = −0.10 to 0.03).

**Fig. 4.**
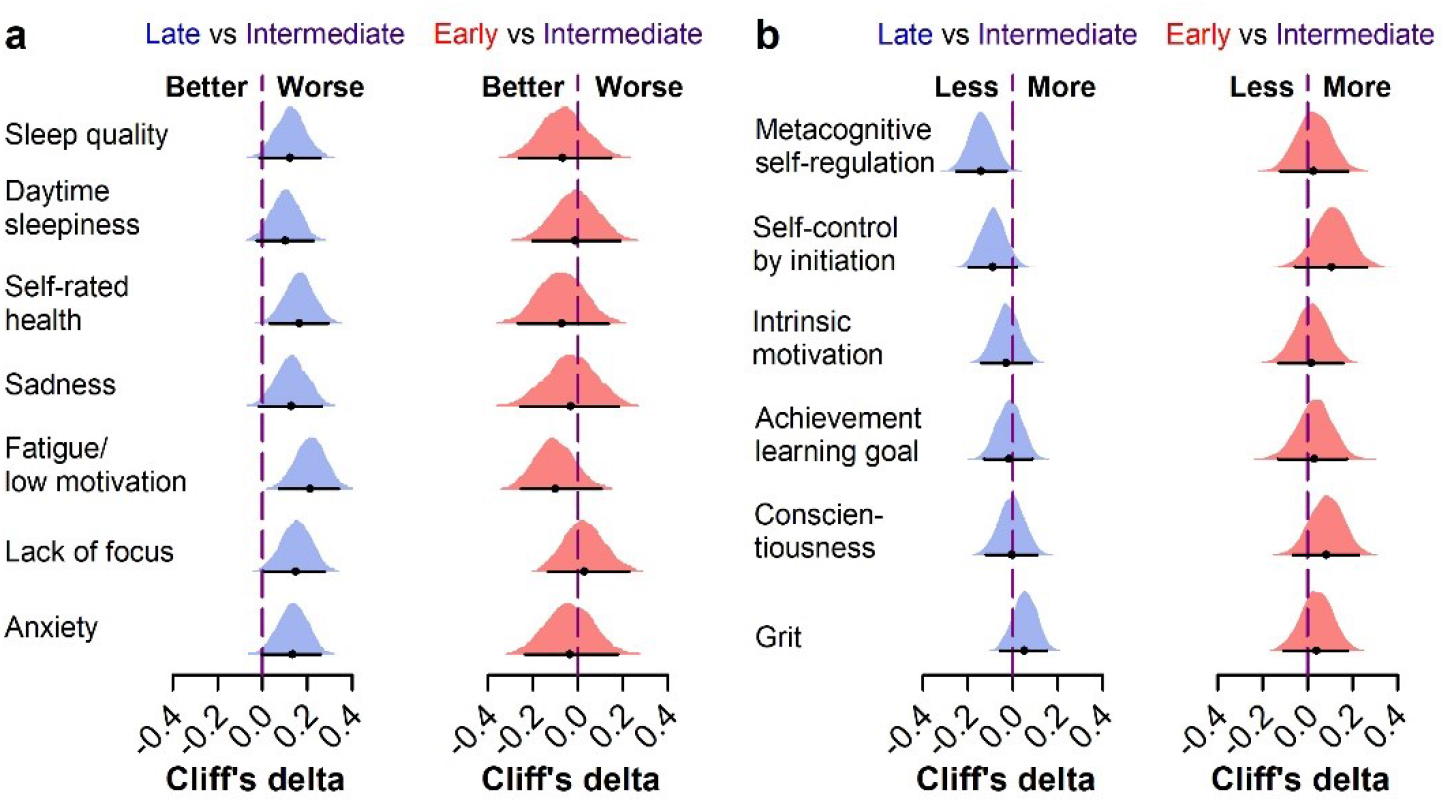
Late-type students had poorer well-being and self-regulation. Well-being measures and psychological correlates of learning were compared between students who were categorized into chronotype groups using their Learning Management System login data. Separate surveys were used to assess (**a**) well-being of students during the school term (Chronotype: early, *n* = 24; intermediate, *n* = 252; late, *n* = 81), and (**b**) psychological characteristics of incoming first-year students (Chronotype: early, *n* = 75; intermediate, *n* = 523; late, *n* = 154). Cliff’s delta was used to estimate effect sizes for early-type and late-type groups relative to the intermediate-type group. The 95% CI and bootstrap sampling distribution is shown for each point estimate of effect size.

Next, we investigated associations of chronotype with psychological correlates of learning (Fig. 4b). Survey data were analyzed for 752 students entering their first year of university whose chronotype was later derived from their LMS logins (early, *n* = 75; intermediate, *n* = 523; late, *n* = 154). Chronotype was inversely associated with metacognitive self-regulation, i.e., purposeful direction of learning through planning, monitoring, and evaluating progress (Kruskal-Wallis test: *H* = 6.8, *P* = 0.034). Multiple comparison tests showed that metacognitive self-regulation was significantly lower in late-type students compared with intermediate-type students (Dunn’s test, *P* = 0.039; Cliff’s delta = −0.14, 95% CI: −0.25 to −0.03), but there was no difference between early-type students and intermediate-type students (Dunn’s test, *P* = 0.73; Cliff’s delta = 0.03, 95% CI: −0.12 to 0.18). Kruskal-Wallis tests did not detect differences between chronotype groups for self-control, intrinsic motivation, learning goals, conscientiousness, or grit (*H* < 5.5, *P* > 0.06 for all measures), and effect sizes were small or negligible with 95% CIs that overlapped zero (Figure 4b; Supplementary Tables 7 and 8).

### Chronotype and lecture attendance

Students’ attendance rates for 337 large lecture courses (≥100 students enrolled in the course) were estimated using time and location data from their Wi-Fi connection logs (38). Wi-Fi confirmed attendance rates for different class start times (08:00, 09:00, 10:00, 12:00, 14:00, and 16:00) were compared between students categorized by their LMS-derived chronotype (early, *n* = 1,563; intermediate, *n* = 12,559; late, *n* = 3,234) (Fig. 5a), adjusting for sex, age, ethnicity, country of citizenship/residency, year of matriculation, school of enrollment, and type of residence. There was a significant interaction between chronotype and class start time on Wi-Fi confirmed attendance (*F*_10, 29302_ = 5.2, *P* < 0.001). Main effects showed that later chronotype (*F*_2, 20362_ = 66.7, *P* < 0.001) and earlier class start times (*F*_5, 29197_ = 88.1, *P* < 0.001) were associated with lower attendance. Overall, Wi-Fi confirmed attendance rates were about 5 percentage points higher in early-type students and about 5 percentage points lower in late-type students, as compared with intermediate-type students. Multiple comparison tests showed that early-type students had higher attendance rates than intermediate-type students for classes that started at 09:00 and 10:00 (Tukey’s test, *P* < 0.017 for both comparisons). Conversely, late-type students had lower attendance rates than intermediate-type students across all class start times (Tukey’s test, *P* < 0.017 for all pairwise comparisons). For comparisons that reached statistical significance, effect sizes were small for early-type students (Cohen’s d range = 0.16 to 0.21) and late-type students (Cohen’s d range = −0.18 to −0.10) (Fig. 5b).

**Fig. 5.**
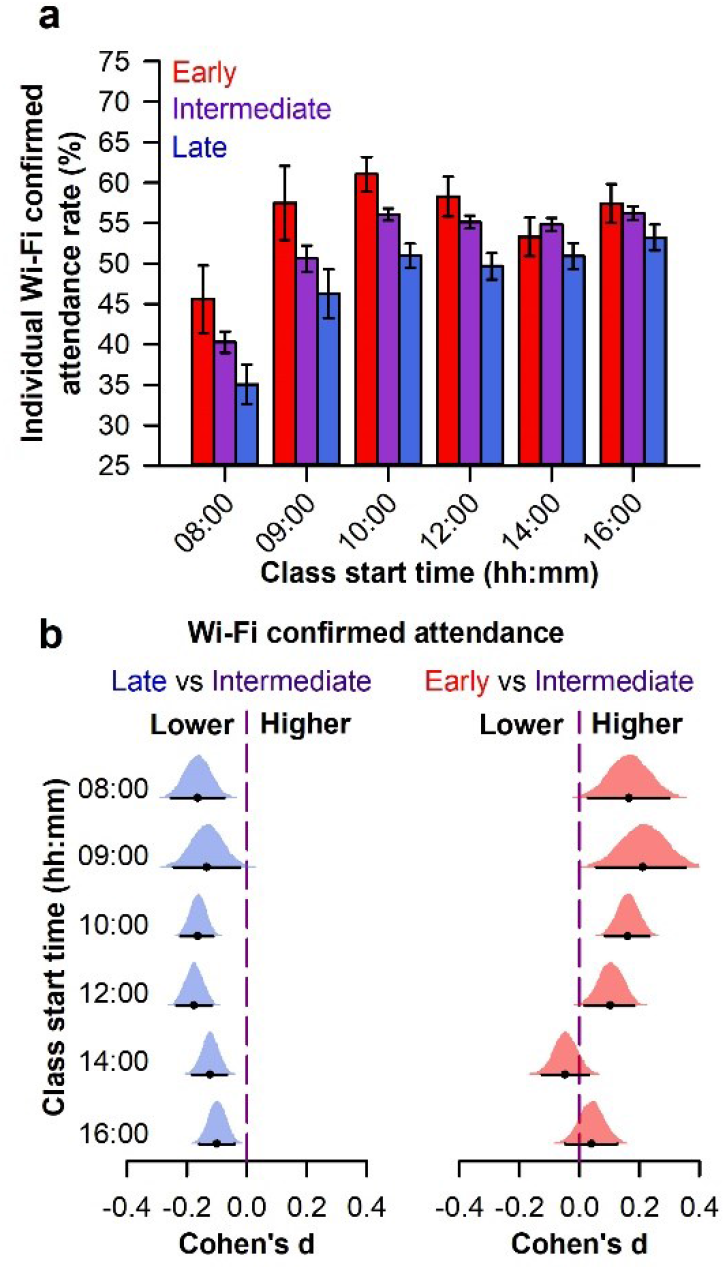
Late-type students had lower Wi-Fi confirmed lecture attendance. Individually determined lecture attendance rates were estimated using students’ Wi-Fi connection logs. Wi-Fi confirmed attendance was compared between students who were categorized into chronotype groups using their Learning Management System login data (early, *n* = 1,563; intermediate, *n* = 12,559; late, *n* = 3,234). (**a**) Across class start times, late-type students had lower Wi-Fi confirmed attendance compared with their peers. Early-type students also had higher attendance rates for morning classes. The mean ± 95% CI is shown. (**b**) Cohen’s d was used to estimate effect sizes for early-type and late-type groups relative to the intermediate group. The 95% CI and bootstrap sampling distribution is shown for each point estimate of effect size.

## Discussion

Our study showed that late-type students have a learning disadvantage relative to their peers. “Burning the midnight oil” is often thought to reflect industriousness that would lead one to eventual success. Our findings turn that common assumption on its head by showing that, in reality, the reverse is true. University-wide chronotyping using LMS login data revealed that late-type students had lower grades for courses held at all times of day. Contrary to the widely held view that shifting courses to the afternoon can remedy this performance gap, late-type students had a lower grade point average during semesters when they had no morning classes. We showed that LMS-derived chronotype was associated with several factors important for academic achievement. Late-type students had shorter nocturnal sleep on days with morning classes, poorer self-rated health, greater frequency of poor mood, lower metacognitive self-regulation, and lower lecture attendance. Our findings suggest that multiple factors converge to impair learning in late-type students.

Our results confirm and extend prior work demonstrating that LMS-derived chronotype is associated with grade point average. We found that late-type students at a large university in Singapore (*n* = 33,645) had lower grades than their peers for courses held in the morning and afternoon/evening, hence reproducing findings from a large university in the midwestern United States (*n* = 14,894) (37). We also showed that the academic disadvantage in late-type students was present for all individual class start times and did not depend on the number of days per week that students had morning classes. Notably, grade point average decreased in all chronotype groups with more days of morning classes, suggesting that even early-type students may benefit from fewer morning classes. The cumulative grade point average in late-type students was about a quarter of a standard deviation lower compared with intermediate-type students (Cohen’s d = −0.27). This effect size is substantively important in the context of academic achievement (39, 40), as it represents a ∼10-point difference in percentile rank. In our study, performance of the average student in the late-type group corresponded to the 39^th^ percentile of performance in the intermediate-type group. Interventions with a comparable effect size on students’ performance are given serious consideration by educational decision-makers. Hence, the academic disadvantage in late-type students is important to address.

Previous studies that measured chronotype based on self-report have produced mixed findings on the relationship between chronotype and grades for morning versus afternoon courses (30-32, 34, 35). Only one study in university students demonstrated a significant interaction between chronotype and time of day on grade point average, in which high morningness was associated with better grades for courses that started at 08:00 or 08:30, but not for later start times (31). Consistent with this finding, a study that compared test scores for a course offered at 08:00 or 14:00 found that later chronotype was associated with lower performance in the morning but not in the afternoon (32). These findings could be explained by greater circadian desynchrony and shorter sleep in late-type students for morning classes. However, other studies found that late-type students had lower grades than early-type students in both morning classes and afternoon classes (34, 35), which is consistent with large-scale data using the LMS chronotyping method (37) including the present study. Hence, there is growing evidence that late-type students show worse performance than their peers even when their classes are scheduled later in the day.

We showed that the LMS chronotyping method can be used to categorize students into groups that differ in their diurnal sleep-wake behavior. On non-school days, sleep onset and sleep offset were later in late-type students and earlier in early-type students, but there were no group differences in nocturnal sleep duration. In contrast, late-type students obtained shorter sleep than their peers on school days due to going to bed late and waking up earlier than usual. Our findings are consistent with previous studies in university students demonstrating that morning-type students obtain more sleep than evening-type students on weekdays (6, 41–43). We extended prior work by demonstrating that nocturnal total sleep time was shorter (by about 40 minutes) in late-type students only when their first class of the day took place in the morning. Comparable results were obtained in adolescents attending schools with morning and afternoon shifts, in which late-type students had shorter sleep than early-type students for the morning shift but not the afternoon shift (7, 8). These studies provide evidence that morning classes contribute to chronic sleep restriction in late-type students, which could contribute to daytime sleepiness and lower grades. In our study, however, the association between chronotype and grade point average did not vary by the numbers of days per week that students had morning classes, and late-type students performed more poorly even when they did not have morning classes. It is therefore unlikely that short sleep and circadian desynchrony can fully explain lower academic achievement in late-type students.

We found that self-rated measures of health and mood tended to be worse in late-type students. Previous studies showed that eveningness was associated with negative mental and physical health outcomes, including greater severity of depression symptoms (18, 44), mood disorders and emotional problems (17), higher BMI or overweight/obesity (45, 46) and metabolic dysregulation (47, 48). Eveningness is also more prevalent among patients with mood disorders, substance abuse, and sleep disorders (17, 49). Although we did not investigate specific health outcomes, our results are consistent with other studies demonstrating that later chronotype was associated with worse self-rated health (13, 14). There was a small effect of chronotype on mood, in which late-type students reported a greater frequency of feeling fatigue / low motivation and lack of focus. However, we did not detect differences in sleep quality or daytime sleepiness between chronotype groups. The link between chronotype and health/mood could be explained by several mechanisms including circadian desynchrony, inadequate sleep, personality traits, and psychosocial characteristics (50).

We found that students with a late diurnal rhythm of LMS logins had lower scores for metacognitive self-regulation. Our results extend previous studies in adolescents showing that self-reported eveningness was associated with lower self-regulation (19, 20). Important aspects of self-regulation include monitoring, evaluating, and managing one’s goal-directed activities (as opposed to merely having knowledge). Late-type students may give less forethought to their learning strategies, or they may be less inclined to correct behavior that they know undermines their best interests. This has implications for both learning and sleep behavior because poor self-regulation skills may hinder turning intentions into actions, leading to goal-inconsistent behaviors such as procrastination. A recent study in university students found that chronotype partially mediated the effect of self-regulation on bedtime procrastination (51), i.e. failing to go to bed at the intended time despite the opportunity to do so (52). Therefore, poorer self-regulation skills in late-type students may contribute to late bedtimes and short nocturnal sleep on school nights. In contrast to previous studies (21–29), chronotype was not associated with self-control, intrinsic motivation, achievement learning goals, conscientiousness, and grit. However, earlier work defined chronotype based on diurnal preference (i.e. morningness or eveningness), which represents a psychological construct that may be more strongly associated with psychological correlates of learning, as compared with an objective measure of diurnal behavior (i.e., LMS logins). Hence, associations between chronotype and psychological correlates of learning may depend strongly on how chronotype is conceptualized and defined.

We provided the first objective evidence that late-type students have lower class attendance than their peers for both morning and afternoon classes. Early-type students also had higher attendance rates for morning classes than the other chronotype groups. These results support the widely held assumption that early-type students are better suited for morning classes, but they also suggest that late-type students are less likely to attend lectures even when they are scheduled at a more favorable circadian phase. Lecture attendance was also lower in all chronotype groups for classes at 08:00, which may indicate that this is too early in the morning for all students. Our findings extend survey studies in university students that assessed self-reported attendance without considering time of day that classes were held. Similar to our results, evening-type students reported missing more classes than morning-type students (6, 41). A study of high school students that used attendance data from the school registration system also found that later chronotype was associated with more days of sick leave (53). In our study, reasons for lower attendance in late-type students could be related to multiple factors including oversleeping, feeling tired, poorer health (e.g. falling sick), lower mood, or lower self-regulation.

Our study showed that several factors may contribute to poorer learning in late-type students. Short nocturnal sleep (54, 55), disturbed mood (56, 57), lower metacognitive self-regulation (58), and absenteeism (59, 60) have all been linked to lower academic performance. We did not attempt to model the relative contributions of these factors to academic achievement because each factor was examined in a different subpopulation. Even if all data were available in the same students, it would be challenging to determine which factors mediate or moderate effects of other variables, and the direction of effects. Later chronotype may impair performance through effects of short sleep and circadian desynchrony. Conversely, students who have lower grades may feel pressured to work later into the night to keep pace with their peers. Likewise, skipping classes could result in students displacing sleep to catch up with their work. Hence, there may be reciprocal relations between chronotype and academic achievement (3). Additionally, lower grades could contribute to poorer mood and well-being, especially if students base their self-worth on academic achievement. Inadequate sleep and mood disturbances also have a bidirectional relationship (61) and could influence students’ self-regulation and attendance. Finally, it should be highlighted that students’ daily schedules may influence their chronotype (9). In summary, it would be difficult to construct a theoretical model that reliably explains the relationship between chronotype and academic achievement. Nonetheless, LMS-derived chronotype can be used by universities and researchers as a marker that is associated with learning-related behaviors and characteristics.

The LMS chronotyping method has several strengths and weaknesses. In contrast to self-report instruments for assessing chronotype, the LMS chronotyping method provides an objective readout of students’ diurnal behavior that associates with sleep. However, the times that students log in to the LMS may be influenced by environmental factors including their social activities, work and extracurricular commitments, and family/living situation. A student’s diurnal rhythm of LMS logins does not necessarily reflect his/her sleep-wake pattern, but it provides confirmation of when he/she is awake and interacting with the LMS. Another drawback is that LMS data cannot be used to estimate daily changes in sleep behavior because students typically interact with the LMS only a few times each day. Rather, the login data accumulate over time, providing an aggregate view of each student’s diurnal activity rhythm. Nonetheless, an important advantage of the LMS chronotyping method over survey methods is that it allows for large-scale (e.g., university-wide) determination of chronotype using passively collected data. This eliminates the need to recruit students to assess their chronotype and reduces sources of bias associated with survey studies (response bias, self-selection bias, self-reporting bias). This method enabled us to investigate associations of chronotype and class start times with grades and lecture attendance at an unprecedented scale.

Insights from our study can be used by universities to improve learning. Building on prior work, we showed that early morning classes are bad for students’ academic performance, sleep, and lecture attendance (38). The negative effect of early classes was greatest in late-type students who went to bed later and slept less than their peers. Delaying the start of school would improve sleep and circadian synchrony in most students, while providing the greatest benefit for late-type students. Universities could also use the LMS chronotyping method to match students by their chronotype to class start times and examination times that are most favorable, especially for courses that are offered at multiple times of day. Given that chronotype-based shift scheduling improves self-reported sleep duration and well-being in shift workers (62), a similar approach could be used by universities to reduce the frequency of morning classes in late-type students. In addition, the LMS chronotyping method can potentially be used to identify students who would benefit from counseling related to sleep behavior, mental health, or strategies for improving metacognition and self-regulation skills (63). These intervention strategies may help to close the performance gap in late-type students, especially if classes and examinations are also shifted later. In conclusion, our study demonstrates that students’ diurnal interactions with the LMS provide valuable information on their sleep behavior and learning. In future work, data collected on these platforms can be used to investigate the factors that drive students’ diurnal behavior and the consequences for learning and health.

## Materials and Methods

### Ethics Statement

Permission to analyze university-archived student data was obtained from the NUS Institute for Applied Learning Sciences and Educational Technology (ALSET). ALSET stores and links de-identified data for educational analytics research, including demographic information (age, sex, ethnicity, country of citizenship/residency, academic year of matriculation, school of enrolment within the university, and type of residence), course enrolment and grades, interactions with the LMS, and Wi-Fi connection logs. Students consented to making their data available for research when they enrolled at NUS and signed the Student Data Protection Policy. Analyses of university-archived data were exempt from review by the NUS Institutional Review Board (IRB) because they were performed retrospectively on data that were de-identified to the researchers. The research was approved by the NUS Learning Analytics Committee on Ethics (LACE) which oversees educational research. Students who took part in surveys and actigraphy studies provided written informed consent, and the procedures were approved by the NUS IRB and LACE.

### Learning Management System (LMS)-derived chronotype

Students’ logins on the LMS were analyzed over 5 semesters (from 2016/17 semester 2 to 2018/19 semester 2). The starting dataset comprised 18.9 million logins from 40,511 students. The dataset included all undergraduate students who enrolled in at least one course. Students from the Faculty of Medicine and Faculty of Dentistry were excluded because these schools have a different type of curriculum and grading system than the other schools at NUS. Login data were sorted by students’ course timetables into non-school days (days with no classes) and school days (days with classes). Data were extracted for 33,645 students who had at least 24 logins on non-school days and at least 24 logins on school days. This criterion ensured that there were sufficient data for constructing individual diurnal profiles of login activity. As described in prior research (37), each student’s time-stamped logins were analyzed using circular statistics to determine the median login phase on non-school days. The population distribution of median login phase was then used to assign participants to chronotype categories. Early-type students (*n* = 3,965) and late-type students (*n* = 5,893) were defined as having a median login phase that was more than one standard deviation earlier or later than the population median, respectively. Intermediate-type students (*n* = 23,787) had a median login phase that was within one standard deviation of the population median. Social jet lag in each student was calculated as the difference in login phase on non-school days relative to school days (i.e., positive values indicated a later phase on non-school days, and negative values indicated an earlier phase on non-school days). Circular statistics were calculated with the “circular” package (version 0.4-93) using R statistical software (version 3.6.3) (64).

### Academic performance

Students’ course grades were extracted for the 5 semesters that LMS data were analyzed. At NUS, students are given a letter grade that is converted to a number for calculating the grade point (A+ = 5.0, A = 5.0, A-= 4.5, B+ = 4.0, B = 3.5, B-= 3.0, C+ = 2.5, C = 2.0, D+ = 1.5, D = 1.0, F = 0.0). Students earn course credits based on the estimated workload hours per week, and the grade point average represents the cumulative performance weighted by the credits earned in each course. ANOVA was used to test for differences in grade point average between chronotype groups (early, *n* = 3,965; intermediate, *n* = 23,787; late, *n* = 5,893).

In other analyses, we investigated whether differences in academic performance between chronotype groups varied by time of day. At NUS, it is common for an individual course to have multiple class start times (e.g., a lecture on Tuesday at 10:00 and a small tutorial class on Thursday at 14:00). We therefore analyzed the data using different approaches to address heterogeneity of students’ timetables. In the first analysis, course grades were sorted by the principal meeting time of the course (e.g., lecture or seminar for all enrolled students) without considering the timing of smaller class meetings. In the second analysis, data were sorted into morning and afternoon/evening courses based on the meeting times of all classes within each course. Morning courses were defined as having all classes (e.g., lectures, tutorials, and laboratories) start before 12:00, and afternoon/evening courses were defined as having all classes start at 12:00 or later. The remaining courses had class meeting times that were held in both the morning and the afternoon/evening. In the third analysis, data were sorted in each semester by the number of days per week that students had morning classes. The grade point average was calculated using all grades that a student obtained in the semester irrespective of times that classes were scheduled. The second and third analyses were restricted to students who earned 20 course credits during the semester (the mode of the distribution for course credits) to ensure that they had a comparable total workload. Separate linear mixed models were used to test the interaction of chronotype with class start time (Analysis 1: 08:00, 09:00, 10:00, 11:00, 12:00, 13:00, 14:00, 15:00, 16:00, 17:00, 18:00), course start time (Analysis 2: morning only, afternoon only, morning and afternoon), and days of the week with morning classes (Analysis 3: 0, 1, 2, 3, 4, 5) on grade point average.

### Sleep behavior

Sleep behavior in NUS undergraduates was assessed during the school term using surveys and actigraphy recordings. Data from 2 studies were combined and sorted by students’ LMS-derived chronotype (**Supplementary Methods**). The dataset comprised 357 students with sleep survey data (Chronotype: early, *n* = 24; intermediate, *n* = 252; late, *n* = 81), including 261 students with 10-42 days of actigraphy data (Chronotype: early, *n* = 18; intermediate, *n* = 189; late, *n* = 54). The sleep survey was used to assess students’ self-reported bedtime, wake-up time, and nocturnal sleep duration on non-school days and school days. The midpoint of TIB was calculated from each student’s bedtime and wake-up time. The survey also assessed various aspects of well-being including sleep quality, daytime functioning, health, and mood using multiple choice questions (**Supplementary Methods**). Objective sleep data were analyzed in participants who wore an actigraphy watch (Actiwatch Spectrum Plus or Actiwatch 2; Philips Respironics Inc., Pittsburgh, PA) on their non-dominant hand for either 6 weeks (study 1, *n* = 148) or 2 weeks (study 2, *n* = 115), including 2 students who took part in both studies. Daily sleep diaries and Actiwatch event markers were used to determine the time-in-bed intervals (**Supplementary Methods**), and sleep scoring was performed using Actiware software (version 6.0.9). In each student, actigraphy-estimated sleep variables (sleep onset, sleep offset, midpoint of sleep, nocturnal total sleep time) were determined for each nocturnal sleep recording. Data were sorted by non-school days and school days, and linear mixed models were used to test for interactive effects of chronotype and type of day (school day, non-school day) on each sleep variable. Secondary analyses were performed to compare sleep behavior between chronotype groups after sorting students’ data by their first class of the day. Data were binned by morning and afternoon/evening classes because there was insufficient data across chronotype groups for sorting by individual class start times.

### Psychological correlates of learning

Students’ psychological characteristics were assessed in a survey administered to incoming first-year students at NUS. The survey comprised about 20 instruments/scales for assessing non-cognitive constructs that are relevant for students’ learning. We limited our analysis to constructs that have been shown to associate with students’ morningness/eveningness preference (19–29), including self-regulation, self-control, intrinsic motivation, achievement learning goals, conscientiousness, and grit (**Supplementary Methods**). The survey was offered to all students who enrolled in the first semester of the 2018/19 academic year, and it was completed prior to the start of classes in August, 2018. Among the 897 first-year students who took part in the survey, there were 752 participants who had sufficient LMS login data in their first year of university for chronotype categorization (early, *n* = 75; intermediate, *n* = 523; late, *n* = 154). Kruskal-Wallis tests were used to test for differences in scores for psychological instruments between chronotype groups.

### Wi-Fi confirmed attendance

Students’ de-identified Wi-Fi connection logs were used to determine the time and location that their Wi-Fi enabled devices associated with the NUS wireless network (65) (**Supplementary Methods**). Students were confirmed as present for a lecture if they connected to a Wi-Fi access point in their lecture hall during class hours. Previously, we showed that Wi-Fi confirmed attendance correlated strongly with instructor-reported attendance (38). We investigated Wi-Fi confirmed attendance for 337 lecture courses with enrollment of >100 students, as described in our earlier work (38). In each course, individually determined Wi-Fi attendance rates were calculated by dividing the number of lectures in which a student was detected by the total number of lectures held in the semester. Among the 23,391 unique students who were enrolled in these courses, there were 17,356 students who were categorized by their LMS-derived chronotype (early, *n* = 1,563; intermediate, *n* = 12,559; late, *n* = 3,234). A linear mixed model was used to test the interaction of chronotype with class start time (08:00, 09:00, 10:00, 12:00, 14:00, 16:00) on Wi-Fi confirmed attendance.

### Statistical comparisons

We implemented two statistical frameworks in each analysis. First, we performed significance-based hypothesis testing to test for differences in behavior between chronotype groups. Second, we implemented estimation statistics to provide a point estimate of effect size and its confidence interval. The latter approach was used for comparing the effect of being an early-type or late-type on behavior, expressed relative to the intermediate-type group that served as the reference.

One-way ANOVA was used to test for differences between chronotype groups in overall grade point average and social jet lag. Multiple comparisons were performed using Tukey’s test (threshold for significance, *P* < 0.05). Effect size was estimated using Cohen’s d. Kruskal-Wallis tests were used to test for differences between chronotype groups for ordinal data (e.g., sleep quality, mood, and psychological correlates of learning). Multiple comparisons were performed using Dunn’s test (threshold for significance, *P* < 0.05). Effect sizes were estimated using Cliff’s delta.

Linear mixed models were used to test for differences between chronotype groups for grade point average, sleep, and Wi-Fi confirmed attendance across time points/periods. Each model included chronotype as a fixed factor and student as a random factor. The repeated fixed factor was a time-related variable (e.g., class start time, type of day, or time of day). Covariates in each model included sex (female, male), age in years as a continuous variable, ethnicity (Chinese, Indian, Malay, Others), country of citizenship/residency (Singaporean, Singapore Permanent Resident, Foreign), academic year of matriculation (2011/12 or earlier, 2012/13, 2013/14, 2014/15, 2015/16, 2016/17, 2017/18, 2018/19), school of enrollment (Faculty of Arts & Social Sciences, Faculty of Engineering, Faculty of Science, Business School, School of Design and Environment, School of Computing, Faculty of Law, Yale-NUS College, Yong Loo Lin School of Medicine Nursing Program, School of Continuing and Lifelong Education, Yong Siew Toh Conservatory of Music, Multi-disciplinary Degree Programs, or Others which comprised students enrolled at multiple schools or who transferred between degree programs), and type of residence (on campus, off campus, mixed). The reference group was intermediate chronotype, female, Chinese, Singaporean, 2017/18 academic year of matriculation, Faculty of Arts & Social Sciences, and off campus residence. Linear mixed models for sleep variables included the same covariates and reference categories, except that class year (year 1, year 2, year 3, year 4) was used instead of the academic year of matriculation, and year 2 was the reference. An F-test was used to test for fixed effects (threshold for statistical significance, *P* < 0.05), with degrees of freedom estimated using Satterthwaite’s method. If there was a significant interaction between chronotype and time/day, multiple comparisons were performed between chronotype groups at different time points using Tukey’s test. If the interaction term was not significant, we examined main effects of chronotype and time. Linear mixed models were implemented using the “lmerTest” package (version 3.1-3) in R Statistical Software (version 4.1.0) (66). Tukey tests were performed using the “emmeans” package (version 1.6.1). Effect sizes of chronotype on grade point average, sleep variables, and Wi-Fi confirmed attendance were measured using Cohen’s d. Effect sizes were calculated with the “dabest” package (version 0.3.0) using Python 3.7.8 (67).

## Acknowledgments and funding sources

We thank researchers and students in the Chronobiology and Sleep Laboratory for collecting data; researchers and administrative staff at the NUS Institute for Applied Learning Sciences and Educational Technology (ALSET) and NUS Information Technology (NUS IT) for supporting analyses of university-archived data; and Dr. Robert K. Kamei for his contributions to ALSET and surveys used in the research. Data storage and management were supported by the NUS Office of the Senior Deputy President & Provost and ALSET. The work was funded by the Ministry of Education, Singapore (MOE2019-T2-2-074), the National Research Foundation, Singapore (NRF2016-SOL002-001), and the Université Sorbonne Paris Cité (USPC)-National University of Singapore (NUS) Joint Innovative Projects in Higher Education (2018-01-EDU/USPC-NUS). The funders had no role in conducting the research.

## Data availability

The sleep survey data and actigraphy data that support the findings of this study are available on request from the corresponding author. University-archived data cannot be shared publicly because of legal and university restrictions where the research was conducted. In compliance with the Singapore Personal Data Protection Act, data stored on the NUS Institute for Applied Learning Sciences and Educational Technology (ALSET) Data Lake is defined as personal data and cannot be shared publicly without student consent. Data can be accessed and analyzed on the ALSET Data Lake server with approval by the NUS Learning Analytics Committee on Ethics, in accordance with NUS data management policies. Researchers who wish to access the data should contact ALSET at NUS.

## Supplementary Methods

### Sleep & actigraphy studies

Sleep behavior was assessed by combining data for 2 studies that implemented similar research procedures. At the start of each study, students completed an online survey of their sleep habits and well-being. Participants then wore an actigraphy watch to record their natural sleep behavior during the school semester. The first study recruited students aged 18-25 years to take part in a 6-week research study (38). Participants were non-smokers in good general health with a body mass index between 18.5-27.0 kg/m^2^. There were 202 students who enrolled in the study and completed the survey of sleep habits and well-being. Among these students, there were 20 individuals who were excluded from actigraphy analyses because of missing or incomplete data (withdrew from the study, *n* = 13; non-compliance, *n* = 5; poor quality data, *n* = 2). The remaining 182 students had 27-42 days of actigraphy data per individual.

The second study recruited students who were enrolled in a course on strategies for learning better (ALS1010 Learning to Learn Better) which was open to undergraduate students from all disciplines. All students enrolled in the course were eligible and there were no exclusionary criteria. There were 216 students who completed the survey of sleep habits and well-being, including 198 students who also agreed to take part in a 2-week actigraphy study. Actigraphy data were excluded for 73 students with missing or poor-quality data (withdrew from study, *n* = 4; non-compliance, *n* = 63; technical problems with the actigraphy watches, *n* = 6). The remaining 125 participants had 10-14 days of actigraphy data per individual. In these participants, survey data were excluded for 3 students who had already participated in the first sleep/actigraphy study.

Across the 2 studies, there were 415 unique students who completed the sleep survey, and 305 unique students who provided 10-42 days of actigraphy data. Analyses were restricted to participants who were assigned to a chronotype category based on their LMS login data. We excluded survey data from 58 students and actigraphy data from 44 students who had insufficient LMS data to derive their chronotype. The final dataset included 357 students with sleep survey data (Chronotype: early, *n* = 24; intermediate, *n* = 252; late, *n* = 81), and 261 students with actigraphy data (Chronotype: early, *n* = 18; intermediate, *n* = 189; late, *n* = 54).

Students in both studies were instructed to wear the actigraphy watch all the time except when taking part in activities that might damage the device. Students pressed an event marker button on their watch when going to bed/ waking up and when putting on/ taking off the actigraphy watch. They also completed a daily diary of times that they slept or removed the actigraphy watch. Actigraphy data were collected in 30-s epochs and analyzed using Actiware software (version 6.0.9). Time-in-bed intervals were marked in the actogram using participants’ event marker presses and sleep diary entries. Sleep scoring was performed using the medium wake-sensitivity threshold (threshold = 40 activity counts) and a 10-minute immobility threshold for determining sleep onset and sleep offset.

### Sleep survey items

Students’ self-reported bedtimes, wake-up times, and nocturnal sleep duration were assessed using free responses. Sleep behavior on non-school days was assessed with the following questions: “What time do you go to bed on a typical free day/ weekend?”, “What time do you wake up on a typical free day/ weekend?”, and “How many hours of sleep do you get at night on a typical free day or weekend?” Sleep behavior on school days was assessed with the following questions: “What time do you go to bed on a typical school day?”, “What time do you wake up on a typical school day?”, and “How many hours of sleep do you get at night on a typical school day?”

Sleep quality was assessed with the following question modified from the Pittsburgh Sleep Quality Index (68): “In the last two weeks, how would you rate your sleep quality?” Response options were “Very good”, “Fairly good”, “Fairly bad”, and “Very bad”. Daytime sleepiness was assessed using the following scenario that was modified from the School Sleep Habits Survey (69): “People sometimes feel sleepy during the daytime. During your daytime activities on a typical school day, how much of a problem is it for you to stay awake? (i.e. feeling sleepy or struggling to stay awake)” Response options were “No problem at all”, “A little problem”, “More than a little problem”, “A big problem”, and “A very big problem”. Self-rated health was assessed with the following question modified from the School Sleep Habits Survey (69), “How would you rate your health compared to other people your age?” Response options were “Poor”, “Fair”, “Good”, and “Excellent”. For each question, responses were coded from 1 to 4, or from 1 to 5, depending on the number of response options. Responses for self-rated health were reverse coded. Therefore, higher scores indicated worse sleep quality, sleepiness, and health.

Mood was assessed using questions that were modified from the Kutcher Adolescent Depression Scale (70, 71). Students were asked to indicate how they felt on average over the past week across different dimensions of mood. Students were presented descriptions of sadness (“Low mood, sadness, feeling blah or down, depressed, just can’t be bothered”), fatigue/ low motivation (“Feeling tired, feeling fatigued, low in energy, hard to get motivated, have to push to get things done, want to rest or lie down a lot”), lack of focus (“Trouble concentrating, can’t keep your mind on schoolwork or work, daydreaming when you should be working, hard to focus when reading, getting “bored” with work or school”), and anxiety (“Feeling worried, nervous, panicky, tense, keyed up, anxious”). Response options included “Hardly ever”, “Some of the time”, “Most of the time”, and “All the time”. Answers were coded from 0 to 3 based on frequency of symptoms. Higher scores indicated poorer mood.

### Instruments for assessing psychological learning characteristics

Metacognitive self-regulation was assessed using the Motivated Strategies for Learning Questionnaire (MSLQ) (72, 73). Students were presented with 12 statements from the subscale “Cognitive and metacognitive strategies: Metacognitive self-regulation” to assess whether they directed their learning by planning, monitoring, and evaluating progress. Response options were presented on a 7-point Likert scale ranging from “Not at all true of me” (1) to “Very true of me” (7). The average score across items was determined in each student. Higher scores indicated higher metacognitive self-regulation.

Initiation of self-control was assessed using items from the Capacity for Self-Control Scale (74). Students were presented with 3 statements in which initiation of a goal-consistent action is required to oppose a pull toward inaction (e.g., due to impulses, habits, desires, requests/demands by others): “I waste a lot of time before getting down to work”, “I waste time on things that don’t really matter, rather than working on things that do”, and “I just can’t seem to get going, even when I have much to do”. Response options were “Strongly disagree”, “Disagree”, “Neutral”, “Agree”, and “Strongly agree”. Items were reverse coded from 1 to 5 and the average score was determined in each student. Higher scores indicated higher self-control by initiation.

Intrinsic motivation was measured using the 17-item Intrinsic Motivation Scale (75). The scale comprises items assessing the desire for challenging tasks (6 items), focus on personal curiosity and interest (6 items), and the desire for independent mastery (5 items). Students rated how well each description applied to them on a 9-point Likert scale ranging from “Never or definitely no” (1) to “Always” (9). The average score across items was determined in each student. Higher scores indicated higher intrinsic motivation.

Students’ learning goals were assessed using items from the Achievement Goal Inventory (76). Students were presented with statements related to acquiring knowledge and skills and seeking challenges: “I strive to constantly learn and improve in my courses”, “In school I am always seeking opportunities to develop new skills and acquire new knowledge”, “In my classes I focus on developing my abilities and acquiring new ones”, “I seek out courses that I will find challenging”, “I really enjoy facing challenges, and I seek out opportunities to do so in my courses”, and “It is very important to me to feel that my coursework offers me real challenges”. Response options were “Strongly disagree”, “Somewhat disagree”, “Neutral”, “Somewhat agree”, and “Strongly agree”. Items were coded from 1 to 5 and the average score was determined in each student. Higher scores indicated higher achievement learning goals.

Conscientiousness was assessed using the Ten-Item Personality Inventory (77). Participants were presented with a list of personality traits and were asked to rate the extent to which they agreed or disagreed that the trait applied to them. Conscientiousness was assessed with the items, “Dependable, self-disciplined” and “Disorganized, careless”. Responses were provided on a 7-point Likert scale with the options “Strongly disagree”, “Disagree”, “Somewhat disagree”, “Neutral”, “Somewhat agree”, “Agree”, and “Strongly agree”. The values for the 2 items were averaged (the second item was reverse-coded), with higher scores indicating higher conscientiousness.

Grit was assessed using the 8-item Short Grit Scale (78). Students were presented with statements relating to perseverance and passion for long-term goals. Students rated whether each statement described them by selecting one of the following response options: “Not like me at all”, “Not much like me”, “Somewhat like me”, “Mostly like me”, and “Very much like me”. Items were coded from 1 to 5 (some items were reverse-coded) and averaged for each participant. Higher scores indicated higher grit.

### Wi-Fi connection data

Students’ Wi-Fi connection logs included the tokenized student identity, the anonymized media access control (MAC) address used to identify the Wi-Fi enabled device, the name and location descriptor of the Wi-Fi access point, and the start and end time of each Wi-Fi connection. Students’ Wi-Fi connection logs were cross-referenced with their class timetables, which made it possible to determine whether they connected to a Wi-Fi router in their classroom during class hours. Wi-Fi confirmed attendance was investigated over 3 semesters using all available data prior to the COVID-19 pandemic (from 2018/19 semester 1 to 2019/20 semester 1). As described in our previous work (38), lecture courses were included in the analysis if they were held once per week and at least 7 times over the 13-week semester. We analyzed courses that lasted 2 h per session and had an enrolment of at least 100 students, which ensured that comparable types of courses were included across different class start times. The final dataset included 337 lecture courses (08:00, 21 courses; 09:00, 18 courses; 10:00, 89 courses; 12:00, 67 courses; 14:00, 72 classes; 16:00, 70 classes). Among the 23,391 unique students who were enrolled in these courses, 17,356 students had sufficient LMS login data for chronotype categorization. The Wi-Fi confirmed attendance rate for each student was determined in each of the 337 courses. This was calculated as the number of lectures in which a student connected to a Wi-Fi router in the classroom, divided by the total number of lectures held during the semester.

## Supplementary Figures

**Supplementary Fig. 1.**
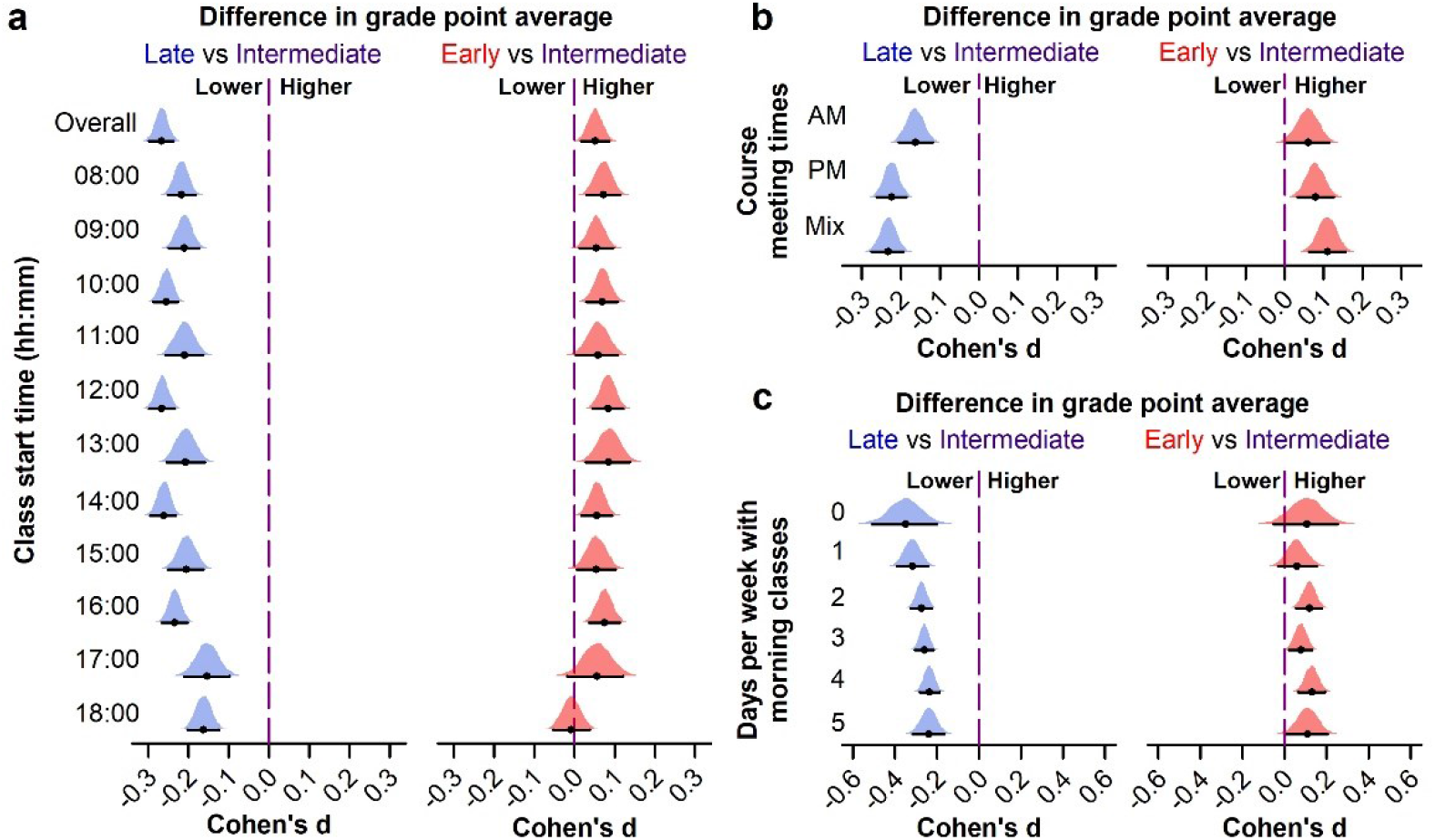
Effect sizes of chronotype on academic performance. Grade point average (GPA) was compared between students who were categorized into chronotype groups (early, *n* = 3,965; intermediate, *n* = 23,787; late, *n* = 5,893) based on their Learning Management System login data. Cohen’s d was used to estimate effect sizes for early-type and late-type groups for (**a**) cumulative GPA and GPA sorted by the primary class start time of a given course (e.g., lecture or seminar), (**b**) GPA sorted by the times of all classes (e.g., lecture, seminar, laboratories, tutorial groups) within a course (AM, all classes started before 12:00; PM, all classes started at 12:00 or later; Mix, classes took place in both the morning and afternoon/evening), and (**c**) GPA sorted by the number of days of the week that students had morning classes. In all analyses, the intermediate-type group was the reference. The 95% CI and bootstrap sampling distribution is shown for each point estimate of effect size.

## Supplementary Tables

**Supplementary Table 1.** Demographic characteristics of students who were categorized by chronotype using their Learning Management System login data.

| | Students<br>enrolled<br>(*n* = 40,511) | Students<br>chronotyped<br>(*n* = 33,645) | Early<br>(*n* = 3,965) | Intermediate<br>(*n* = 23,787) | Late<br>(*n* = 5,893) | *F*/$\chi^2$ | *P* |
| --- | --- | --- | --- | --- | --- | --- | --- |
| **Sex** |  |  |  |  |  |  |  |
| Female | 20,747 (51.2) | 17,657 (52.5) | 2,027 (51.1) | 12,562 (52.8) | 3,068 (52.1) | 4.4 | 0.112 |
| Male | 19,764 (48.8) | 15,988 (47.5) | 1,938 (48.9) | 11,225 (47.2) | 2,825 (47.9) |  |  |
| **Age in years** | 20.8 ± 2.7 | 20.6 ± 2.6 | 21.4 ± 3.3 | 20.6 ± 2.5 | 20.4 ± 2.5 | 231.7 | <0.001 |
| **Ethnicity** |  |  |  |  |  |  |  |
| Chinese | 34,895 (86.1) | 29,262 (87.0) | 3,420 (86.3) | 20,742 (87.2) | 5,100 (86.5) | 40.1 | <0.001 |
| Indian | 1,949 (4.8) | 1,572 (4.7) | 208 (5.2) | 1,151 (4.8) | 213 (3.6) |  |  |
| Malay | 1,273 (3.1) | 1,076 (3.2) | 136 (3.4) | 726 (3.1) | 214 (3.6) |  |  |
| Others | 2,394 (5.9) | 1,735 (5.2) | 201 (5.1) | 1,168 (4.9) | 366 (6.2) |  |  |
| **Citizenship/residency** |  |  |  |  |  |  |  |
| Singapore Citizen | 33,511 (82.7) | 28,484 (84.7) | 3,199 (80.7) | 20,272 (85.2) | 5,013 (85.1) | 71.4 | <0.001 |
| Singapore PR | 2,596 (6.4) | 2,067 (6.1) | 345 (8.7) | 1,407 (5.9) | 315 (5.3) |  |  |
| Foreign | 4,404 (10.9) | 3,094 (9.2) | 421 (10.6) | 2,108 (8.9) | 565 (9.6) |  |  |
| **Academic year of matriculation** |  |  |  |  |  |  |  |
| 2011/12 or earlier | 125 (0.3) | 46 (0.1) | 15 (0.4) | 26 (0.1) | 5 (0.1) | 593.6 | <0.001 |
| 2012/13 | 493 (1.2) | 200 (0.6) | 44 (1.1) | 126 (0.5) | 30 (0.5) |  |  |
| 2013/14 | 5,289 (13.1) | 2,381 (7.1) | 467 (11.8) | 1,547 (6.5) | 367 (6.2) |  |  |
| 2014/15 | 6,617 (16.3) | 5,520 (16.4) | 883 (22.3) | 3,801 (16.0) | 836 (14.2) |  |  |
| 2015/16 | 6,612 (16.3) | 6,245 (18.6) | 806 (20.3) | 4,556 (19.2) | 883 (15.0) |  |  |
| 2016/17 | 6,699 (16.5) | 6,296 (18.7) | 638 (16.1) | 4,603 (19.4) | 1,055 (17.9) |  |  |
| 2017/18 | 7,000 (17.3) | 6,501 (19.3) | 512 (12.9) | 4,731 (19.9) | 1,258 (21.3) |  |  |
| 2018/19 | 7,676 (18.9) | 6,456 (19.2) | 600 (15.1) | 4,397 (18.5) | 1,459 (24.8) |  |  |
| **School of enrollment** |  |  |  |  |  |  |  |
| FASS | 9,029 (22.3) | 7,989 (23.7) | 806 (20.3) | 5,812 (24.4) | 1,371 (23.3) | 392.3 | <0.001 |
| FoE | 8,386 (20.7) | 6,861 (20.4) | 988 (24.9) | 4,690 (19.7) | 1,183 (20.1) |  |  |
| FoS | 6,969 (17.2) | 5,744 (17.1) | 637 (16.1) | 4,033 (17.0) | 1,074 (18.2) |  |  |
| BIZ | 4,110 (10.1) | 3,754 (11.2) | 404 (10.2) | 2,831 (11.9) | 519 (8.8) |  |  |
| SDE | 2,753 (6.8) | 2,356 (7.0) | 230 (5.8) | 1,654 (7.0) | 472 (8.0) |  |  |
| SoC | 2,695 (6.7) | 2,355 (7.0) | 233 (5.9) | 1,654 (7.0) | 468 (7.9) |  |  |
| FoL | 1,198 (3.0) | 957 (2.8) | 140 (3.5) | 681 (2.9) | 136 (2.3) |  |  |
| YNC | 1,110 (2.7) | 54 (0.2) | 11 (0.3) | 35 (0.1) | 8 (0.1) |  |  |
| YLL (Nursing) | 985 (2.4) | 842 (2.5) | 118 (3.0) | 564 (2.4) | 160 (2.7) |  |  |
| SCALE | 802 (2.0) | 713 (2.1) | 166 (4.2) | 439 (1.8) | 108 (1.8) |  |  |
| MUS | 315 (0.8) | 211 (0.6) | 18 (0.5) | 96 (0.4) | 97 (1.6) |  |  |
| MDP | 729 (1.8) | 624 (1.9) | 60 (1.5) | 450 (1.9) | 114 (1.9) |  |  |
| Other (multiple schools or transfers) | 1,430 (3.5) | 1,185 (3.5) | 154 (3.9) | 848 (3.6) | 183 (3.1) |  |  |
| **Type of residence** |  |  |  |  |  |  |  |
| Off campus | 24,905 (61.5) | 20,068 (59.6) | 2,842 (71.7) | 14,078 (59.2) | 3,148 (53.4) | 426.4 | <0.001 |
| On campus | 7,953 (19.6) | 6,411 (19.1) | 499 (12.6) | 4,405 (18.5) | 1,507 (25.6) |  |  |
| Mixed | 7,653 (18.9) | 7,166 (21.3) | 624 (15.7) | 5,304 (22.3) | 1,238 (21.0) |  |  |

All values in the table indicate the *n* (%) except for age (mean ± SD). PR, permanent resident; FASS, Faculty of Arts & Social Sciences; FoE, Faculty of Engineering; FoS, Faculty of Science; BIZ, Business School; SDE, School of Design and Environment; SoC, School of Computing; FoL, Faculty of Law; YNC, Yale-NUS College; YLL (nursing), Yong Loo Lin School of Medicine, Alice Lee Centre for Nursing Studies; SCALE, School of Continuing and Lifelong Education; MUS, Yong Siew Toh Conservatory of Music; MDP, Multi-disciplinary Degree Programs.

**Supplementary Table 2.** Sleep behavior for different LMS-derived chronotype groups.

| Self-reported sleep | LMS-derived chronotype |  |  |  |
| --- | --- | --- | --- | --- |
|  | All<br>(n = 357) | Early<br>(n = 24) | Intermediate<br>(n = 252) | Late<br>(n = 81) |
| **Sleep on non-school days** |  |  |  |  |
| Bedtime (hh:mm) | 01:39 ± 01:25 | 00:41 ± 01:10 | 01:32 ± 01:20 | 02:21 ± 01:30 |
| Wake-up time (hh:mm) | 09:56 ± 01:37 | 08:51 ± 01:35 | 09:48 ± 01:27 | 10:43 ± 01:50 |
| Midpoint of TIB (hh:mm) | 05:47 ± 01:23 | 04:46 ± 01:16 | 05:39 ± 01:16 | 06:35 ± 01:27 |
| Nocturnal sleep duration (h) | 8.2 ± 1.1 | 8.1 ± 1.2 | 8.2 ± 1.0 | 8.3 ± 1.4 |
| **Sleep on school days** |  |  |  |  |
| Bedtime (hh:mm) | 01:04 ± 01:14 | 00:15 ± 01:03 | 00:55 ± 01:08 | 01:47 ± 01:16 |
| Wake-up time (hh:mm) | 08:11 ± 01:12 | 07:48 ± 01:19 | 08:09 ± 01:09 | 08:24 ± 01:18 |
| Midpoint of TIB (hh:mm) | 04:38 ± 01:03 | 04:01 ± 01:06 | 04:32 ± 00:59 | 05:06 ± 01:07 |
| Nocturnal sleep duration (h) | 6.8 ± 1.1 | 7.0 ± 0.7 | 6.9 ± 1.0 | 6.4 ± 1.2 |
| Actigraphy-estimated sleep | All<br>(n = 261) | Early<br>(n = 18) | Intermediate<br>(n = 189) | Late<br>(n = 54) |
|  | All<br>(n = 261) | Early<br>(n = 18) | Intermediate<br>(n = 189) | Late<br>(n = 54) |
| **Sleep on non-school days** |  |  |  |  |
| Sleep onset (hh:mm) | 02:02 ± 01:17 | 01:21 ± 01:29 | 01:56 ± 01:12 | 02:36 ± 01:18 |
| Sleep offset (hh:mm) | 09:05 ± 01:21 | 08:05 ± 01:14 | 09:06 ± 01:18 | 09:21 ± 01:27 |
| Midpoint of sleep (hh:mm) | 05:33 ± 01:13 | 04:43 ± 01:17 | 05:31 ± 01:09 | 05:58 ± 01:15 |
| Nocturnal total sleep time (h) | 6.3 ± 0.9 | 6.1 ± 0.8 | 6.4 ± 0.9 | 6.1 ± 1.0 |
| **Sleep on school days** |  |  |  |  |
| Sleep onset (hh:mm) | 01:48 ± 01:12 | 00:52 ± 01:26 | 01:45 ± 01:09 | 02:17 ± 01:03 |
| Sleep offset (hh:mm) | 08:11 ± 01:08 | 07:38 ± 01:20 | 08:13 ± 01:06 | 08:16 ± 01:07 |
| Midpoint of sleep (hh:mm) | 05:00 ± 01:03 | 04:15 ± 01:21 | 04:59 ± 01:00 | 05:16 ± 00:59 |
| Nocturnal total sleep time (h) | 5.7 ± 0.9 | 6.1 ± 0.6 | 5.8 ± 0.9 | 5.3 ± 0.9 |

Students were categorized into chronotype groups (early, intermediate, late) using their Learning Management System (LMS) login data. The mean ± SD is shown for all sleep variables. TIB, time in bed.

**Supplementary Table 3.** Statistical comparisons of LMS-derived chronotype (early, intermediate, late) and type of day (non-school day, school day) for different sleep variables.

|  | Chronotype |  | Day |  | Chronotype × Day |  |
| --- | --- | --- | --- | --- | --- | --- |
| **Self-reported sleep** | *F* | *P* | *F* | *P* | *F* | *P* |
| Bedtime (hh:mm) | 21.4 | <0.001 | 35.3 | <0.001 | 0.4 | 0.678 |
| Wake-up time (hh:mm) | 9.3 | <0.001 | 186.2 | <0.001 | 9.5 | <0.001 |
| Midpoint of TIB (hh:mm) | 18.8 | <0.001 | 158.3 | <0.001 | 6.1 | 0.002 |
| Nocturnal sleep duration (h) | 2.5 | 0.087 | 166.6 | <0.001 | 5.6 | 0.004 |
| **Actigraphy-estimated sleep** | *F* | *P* | *F* | *P* | *F* | *P* |
| Sleep onset (hh:mm) | 8.1 | <0.001 | 26.5 | <0.001 | 2.5 | 0.085 |
| Sleep offset (hh:mm) | 4.8 | 0.009 | 59.8 | <0.001 | 3.2 | 0.044 |
| Midpoint of sleep (hh:mm) | 6.5 | 0.002 | 71.9 | <0.001 | 1.9 | 0.158 |
| Nocturnal total sleep time (h) | 4.2 | 0.016 | 23.2 | <0.001 | 3.1 | 0.047 |

Sleep variables were determined for non-school days and school days in students who were categorized into chronotype groups (early, intermediate, late) using their Learning Management System (LMS) login data. Linear mixed models were used to test for interactive effects of chronotype and type of day on sleep. Each model included chronotype as a fixed factor, type of day as a repeated fixed factor, and student as a random factor. Covariates included sex, age, ethnicity, country of citizenship/residency, class year, school of enrollment, and type of residence.

**Supplementary Table 4.** Effect sizes of LMS-derived chronotype for different sleep variables.

| Self-reported sleep | Difference relative to Intermediate-type students |  |  |  |
| --- | --- | --- | --- | --- |
|  | Early-type students |  | Late-type students |  |
|  | Mean difference in hours (95% CI) | Cohen's d (95% CI) | Mean difference in hours (95% CI) | Cohen's d (95% CI) |
| **Sleep on non-school days** |  |  |  |  |
| Bedtime (hh:mm) | -0.84 (-1.34 to -0.38) | -0.64 (-1.01 to -0.28) | 0.82 (0.44 to 1.20) | 0.60 (0.32 to 0.88) |
| Wake-up time (hh:mm) | -0.95 (-1.62 to -0.35) | -0.65 (-1.11 to -0.23) | 0.91 (0.50 to 1.37) | 0.59 (0.31 to 0.88) |
| Midpoint of TIB (hh:mm) | -0.88 (-1.44 to -0.40) | -0.69 (-1.13 to -0.31) | 0.93 (0.58 to 1.30) | 0.71 (0.43 to 0.99) |
| Nocturnal sleep duration (h) | -0.03 (-0.47 to 0.55) | -0.03 (-0.46 to 0.53) | 0.10 (-0.25 to 0.44) | 0.09 (-0.23 to 0.40) |
| **Sleep on school days** |  |  |  |  |
| Bedtime (hh:mm) | -0.68 (-1.08 to -0.23) | -0.61 (-0.97 to -0.20) | 0.86 (0.57 to 1.18) | 0.74 (0.47 to 1.02) |
| Wake-up time (hh:mm) | -0.36 (-1.02 to 0.09) | -0.31 (-0.86 to 0.08) | 0.25 (-0.05 to 0.57) | 0.21 (-0.05 to 0.48) |
| Midpoint of TIB (hh:mm) | -0.52 (-1.00 to -0.12) | -0.52 (-1.00 to -0.10) | 0.57 (0.31 to 0.85) | 0.56 (0.29 to 0.85) |
| Nocturnal sleep duration (h) | 0.12 (-0.24 to 0.38) | 0.13 (-0.25 to 0.39) | -0.47 (-0.78 to -0.19) | -0.44 (-0.74 to -0.17) |
| Actigraphy-estimated sleep | Early-type students |  | Late-type students |  |
|  | Mean difference in hours (95% CI) | Cohen's d (95% CI) | Mean difference in hours (95% CI) | Cohen's d (95% CI) |
|  | Mean difference in hours (95% CI) | Cohen's d (95% CI) | Mean difference in hours (95% CI) | Cohen's d (95% CI) |
| **Sleep on non-school days** |  |  |  |  |
| Sleep onset (hh:mm) | -0.57 (-1.28 to 0.08) | -0.46 (-1.04 to 0.08) | 0.63 (0.25 to 1.02) | 0.52 (0.18 to 0.83) |
| Sleep offset (hh:mm) | -1.02 (-1.69 to -0.49) | -0.77 (-1.26 to -0.36) | 0.25 (-0.16 to 0.67) | 0.18 (-0.12 to 0.49) |
| Midpoint of sleep (hh:mm) | -0.79 (-1.43 to -0.22) | -0.68 (-1.21 to -0.18) | 0.44 (0.08 to 0.82) | 0.37 (0.07 to 0.69) |
| Nocturnal total sleep time (h) | -0.22 (-0.66 to 0.14) | -0.24 (-0.71 to 0.16) | -0.31 (-0.63 to -0.0005) | -0.32 (-0.65 to 0.01) |
| **Sleep on school days** |  |  |  |  |
| Sleep onset (hh:mm) | -0.89 (-1.55 to -0.22) | -0.75 (-1.31 to -0.17) | 0.53 (0.21 to 0.85) | 0.46 (0.17 to 0.77) |
| Sleep offset (hh:mm) | -0.58 (-1.29 to -0.03) | -0.52 (-1.14 to -0.02) | 0.05 (-0.27 to 0.38) | 0.05 (-0.25 to 0.34) |
| Midpoint of sleep (hh:mm) | -0.74 (-1.38 to -0.16) | -0.71 (-1.34 to -0.15) | 0.29 (0.01 to 0.58) | 0.29 (0.00 to 0.59) |
| Nocturnal total sleep time (h) | 0.26 (-0.08 to 0.53) | 0.29 (-0.10 to 0.62) | -0.47 (-0.73 to -0.21) | -0.53 (-0.83 to -0.24) |

Students were categorized into chronotype groups (early, intermediate, late) using their Learning Management System (LMS) login data. Effect sizes were measured relative to the intermediate-type group. TIB, time in bed.

**Supplementary Table 5.**
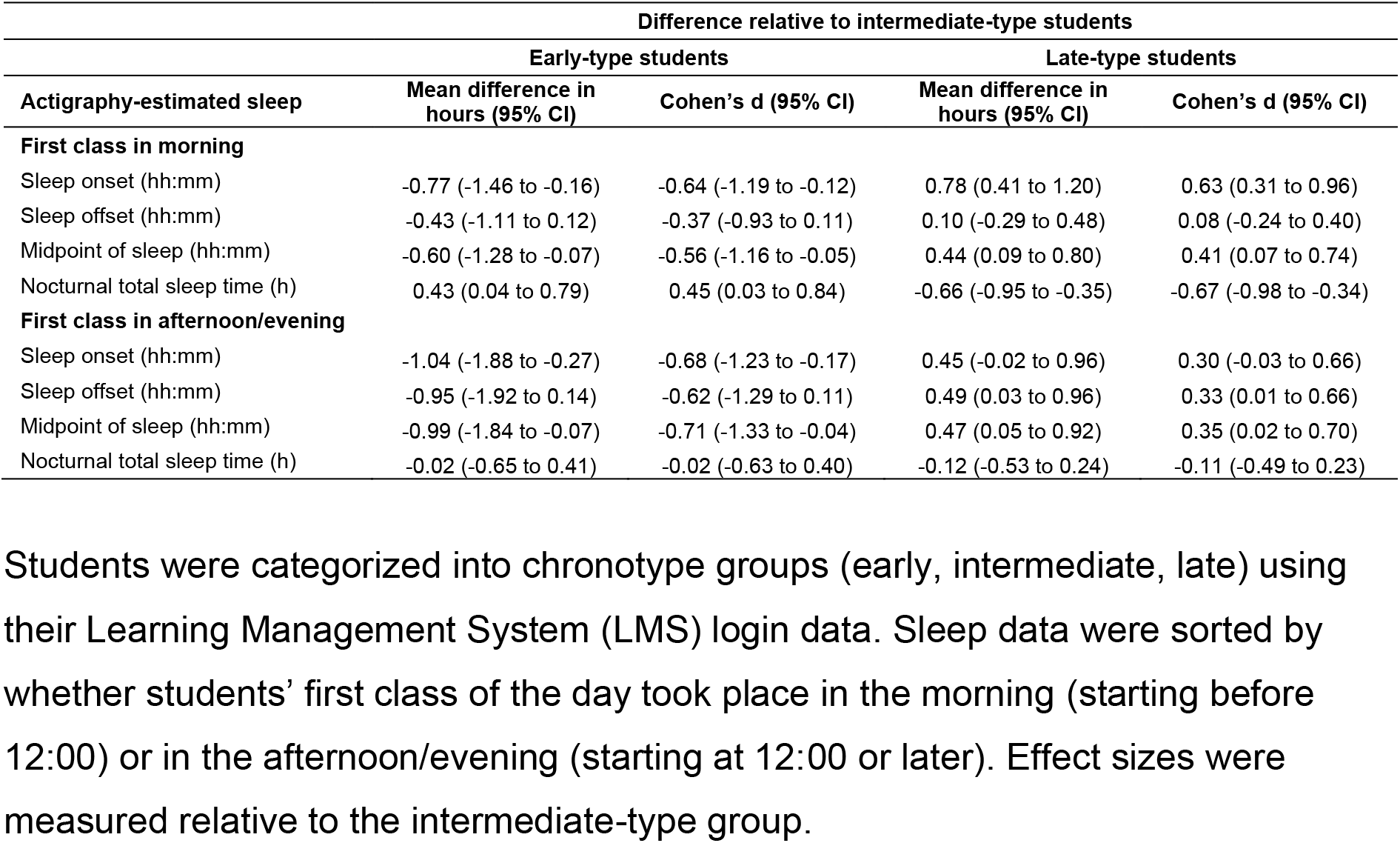
Effect sizes of LMS-derived chronotype for sleep variables sorted by students’ first class of the day.

**Supplementary Table 6.** Well-being measures for different chronotype groups.

| LMS-derived chronotype |  |  |  |  |  |  |
| --- | --- | --- | --- | --- | --- | --- |
| Variable (coded response) | All<br>*n* (%) | Early<br>*n* (%) | Intermediate<br>*n* (%) | Late<br>*n* (%) | *H* | *P* |
| **Sleep quality** |  |  |  |  |  |  |
| Very good (1) | 40 (11.3%) | 3 (13.0%) | 30 (11.9%) | 7 (8.8%) | 4.2 | 0.123 |
| Fairly Good (2) | 205 (57.7%) | 15 (65.2%) | 149 (59.1%) | 41 (51.3%) |  |  |
| Fairly Bad (3) | 94 (26.5%) | 4 (17.4%) | 63 (25.0%) | 27 (33.8%) |  |  |
| Very bad (4) | 16 (4.5%) | 1 (4.3%) | 10 (4.0%) | 5 (6.3%) |  |  |
| **Daytime sleepiness** |  |  |  |  |  |  |
| No problem at all (1) | 34 (9.5%) | 2 (8.3%) | 26 (10.3%) | 6 (7.4%) | 2.5 | 0.281 |
| A little problem (2) | 210 (58.8%) | 15 (62.5%) | 152 (60.3%) | 43 (53.1%) |  |  |
| More than a little problem (3) | 72 (20.2%) | 6 (25.0%) | 44 (17.5%) | 22 (27.2%) |  |  |
| A big problem (4) | 38 (10.6%) | 1 (4.2%) | 28 (11.1%) | 9 (11.1%) |  |  |
| A very big problem (5) | 3 (0.8%) | 0 (0.0%) | 2 (0.8%) | 1 (1.2%) |  |  |
| **Self-rated health** |  |  |  |  |  |  |
| Poor (4) | 5 (1.4%) | 0 (0.0%) | 2 (0.8%) | 3 (3.8%) | 7.1 | 0.029 |
| Fair (3) | 106 (29.8%) | 5 (20.8%) | 71 (28.2%) | 30 (37.5%) |  |  |
| Good (2) | 193 (54.2%) | 15 (62.5%) | 138 (54.8%) | 40 (50.0%) |  |  |
| Excellent (1) | 52 (14.6%) | 4 (16.7%) | 41 (16.3%) | 7 (8.8%) |  |  |
| **Sadness** |  |  |  |  |  |  |
| Hardly ever (0) | 85 (23.9%) | 7 (29.2%) | 62 (24.6%) | 16 (20.3%) | 4.0 | 0.135 |
| Some of the time (1) | 206 (58.0%) | 13 (54.2%) | 151 (59.9%) | 42 (53.2%) |  |  |
| Most of the time (2) | 56 (15.8%) | 4 (16.7%) | 36 (14.3%) | 16 (20.3%) |  |  |
| All the time (3) | 8 (2.3%) | 0 (0.0%) | 3 (1.2%) | 5 (6.3%) |  |  |
| **Fatigue/ low motivation** |  |  |  |  |  |  |
| Hardly ever (0) | 38 (10.7%) | 2 (8.3%) | 30 (11.9%) | 6 (7.5%) | 12.0 | 0.003 |
| Some of the time (1) | 195 (54.8%) | 18 (75.0%) | 143 (56.7%) | 34 (42.5%) |  |  |
| Most of the time (2) | 95 (26.7%) | 3 (12.5%) | 64 (25.4%) | 28 (35.0%) |  |  |
| All the time (3) | 28 (7.9%) | 1 (4.2%) | 15 (6.0%) | 12 (15.0%) |  |  |
| **Lack of focus** |  |  |  |  |  |  |
| Hardly ever (0) | 39 (11.0%) | 1 (4.2%) | 30 (11.9%) | 8 (10.0%) | 5.0 | 0.082 |
| Some of the time (1) | 198 (55.6%) | 16 (66.7%) | 146 (57.9%) | 36 (45.0%) |  |  |
| Most of the time (2) | 96 (27.0%) | 7 (29.2%) | 61 (24.2%) | 28 (35.0%) |  |  |
| All the time (3) | 23 (6.5%) | 0 (0.0%) | 15 (6.0%) | 8 (10.0%) |  |  |
| **Anxiety** |  |  |  |  |  |  |
| Hardly ever (0) | 100 (28.2%) | 7 (29.2%) | 77 (30.6%) | 16 (20.3%) | 4.2 | 0.120 |
| Some of the time (1) | 185 (52.1%) | 14 (58.3%) | 128 (50.8%) | 43 (54.4%) |  |  |
| Most of the time (2) | 56 (15.8%) | 3 (12.5%) | 38 (15.1%) | 15 (19.0%) |  |  |
| All the time (3) | 14 (3.9%) | 0 (0.0%) | 9 (3.6%) | 5 (6.3%) |  |  |

Students were categorized into chronotype groups (early, intermediate, late) using their Learning Management System (LMS) login data. Response options for each item were coded by integers, and non-parametric statistical comparisons between chronotype groups were performed using Kruskal-Wallis tests.

**Supplementary Table 7.** Effect sizes of LMS-derived chronotype for well-being measures and psychological correlates of learning.

|  | **Difference relative to intermediate-type students** |  |
| --- | --- | --- |
|  | **Early-type students**<br>**Cliff's delta (95% CI)** | **Late-type students**<br>**Cliff's delta (95% CI)** |
| **Well-being** |  |  |
| Sleep quality | -0.07 (-0.26 to 0.15) | 0.12 (-0.01 to 0.26) |
| Daytime sleepiness | -0.01 (-0.20 to 0.19) | 0.10 (-0.03 to 0.23) |
| Self-rated health | -0.07 (-0.27 to 0.14) | 0.16 (0.03 to 0.29) |
| Sadness | -0.03 (-0.26 to 0.18) | 0.13 (-0.02 to 0.26) |
| Fatigue/ low motivation | -0.10 (-0.25 to 0.10) | 0.21 (0.07 to 0.34) |
| Lack of focus | 0.03 (-0.13 to 0.23) | 0.15 (0.01 to 0.28) |
| Anxiety | -0.04 (-0.24 to 0.18) | 0.13 (-0.004 to 0.26) |
| **Psychological characteristics** |  |  |
| Metacognitive self-regulation | 0.03 (-0.12 to 0.18) | -0.14 (-0.25 to -0.03) |
| Self-control by initiation | 0.11 (-0.05 to 0.26) | -0.09 (-0.20 to 0.02) |
| Intrinsic motivation | 0.02 (-0.13 to 0.16) | -0.03 (-0.14 to 0.08) |
| Achievement learning goal | 0.03 (-0.13 to 0.18) | -0.02 (-0.13 to 0.09) |
| Conscientiousness | 0.08 (-0.06 to 0.23) | 0.00 (-0.12 to 0.11) |
| Grit | 0.04 (-0.10 to 0.18) | 0.05 (-0.05 to 0.15) |

Students were categorized into chronotype groups (early, intermediate, late) using their Learning Management System (LMS) login data. Survey data for well-being measures were analyzed in 357 students (Chronotype: early, $n = 24$ ; intermediate, $n = 252$ ; late, $n = 81$ ). Survey data for psychological correlates of learning were analyzed in 752 students (Chronotype: early, $n = 75$ ; intermediate, $n = 523$ ; late, $n = 154$ ). Cliff's delta was used to estimate effect sizes for early-type and late-type groups relative to the intermediate-type group.

**Supplementary Table 8.** Statistical comparisons between chronotype groups for psychological correlates of learning.

| Instrument (score range) | LMS-derived chronotype |  |  |  | *H* | *P* |
| --- | --- | --- | --- | --- | --- | --- |
|  | All<br>(*n* = 752) | Early<br>(*n* = 75) | Intermediate<br>(*n* = 523) | Late<br>(*n* = 154) |  |  |
| MSLQ: Metacognitive self-regulation (1-7) | 4.83 (4.25-5.33) | 4.92 (4.25-5.50) | 4.83 (4.33-5.42) | 4.58 (4.17-5.23) | 6.8 | 0.034 |
| CSCS: Self-control by initiation (1-5) | 3.00 (2.33-3.67) | 3.33 (2.17-4.00) | 3.00 (2.33-3.67) | 2.67 (2.33-3.67) | 5.4 | 0.066 |
| Intrinsic Motivation Scale (1-9) | 6.71 (5.88-7.47) | 6.74 (6.18-7.50) | 6.71 (5.88-7.47) | 6.65 (5.82-7.35) | 0.4 | 0.835 |
| AGI: Learning goal (1-5) | 4.17 (3.83-4.67) | 4.17 (3.67-4.67) | 4.17 (3.83-4.50) | 4.17 (3.67-4.50) | 0.3 | 0.851 |
| TIPI: Conscientiousness (1-7) | 5.00 (4.00-5.50) | 5.00 (4.00-6.00) | 5.00 (4.00-5.50) | 4.50 (4.00-5.50) | 1.3 | 0.529 |
| Short Grit Scale (1-5) | 3.38 (3.00-3.75) | 3.38 (3.00-3.75) | 3.38 (3.00-3.75) | 3.50 (3.12-3.75) | 1.0 | 0.608 |

Students were categorized into chronotype groups (early, intermediate, late) based on their Learning Management System (LMS) login data. Kruskal-Wallis tests were used to compare psychological correlates of learning between chronotype groups. Values indicate the median and interquartile range. MSLQ, Motivated Strategies for Learning Questionnaire; CSCS, Capacity for Self-Control Scale; AGI, Achievement Goal Inventory; TIPI, Ten-Item Personality Inventory.

